# Organization of Myosin H in the Apical Complex of *Toxoplasma Gondii* Revealed by 3D Single-Molecule Super-Resolution Microscopy

**DOI:** 10.64898/2026.04.23.720434

**Authors:** Ashwin Balaji, Li-av Segev Zarko, Andrew E.S. Barentine, John C. Boothroyd, W.E. Moerner

## Abstract

*Toxoplasma gondii* is a single-celled eukaryotic parasite with prolific invasion capability. The parasite uses an apical complex comprised of proteinaceous structures and secretory organelles to efficiently enter host cells. As a result, the apical complex remains a vital structure of interest, with many studies dedicated to understanding its protein organization. One such protein is the motor Myosin H (MyoH), which is indispensable for parasite motility and host cell invasion. Given the small size of the complex, roughly a diffraction-limited volume in the visible, high-resolution techniques are required to make precise determinations of protein organization. In this work, we use 3D single-molecule localization microscopy in both traditionally fixed and gel-expanded parasites to localize the indispensable motor Myosin H within the apical complex. Labeling of the N- and C-terminus of MyoH in fixed parasites resolved the orientation of the motor protein in the apical complex, showing the motor head radially exterior to the tail. Two-color imaging of MyoH with tubulin in fixed parasites allowed for localization of the MyoH termini relative to the conoid, a barrel of tubulin-based fibers in the apical complex and showed the MyoH tail toward the interior face of the conoid and the head at the conoid exterior. Gel expansion showed improved labeling density for both tubulin and MyoH but altered MyoH localization, highlighting the nuanced effects of gel expansion on protein organization.

**Statement of Significance:** This work employs 3D single-molecule super-resolution microscopy to provide quantitative physical analysis of the spatial organization of a vital myosin motor, MyoH, in the model apicomplexan parasite *Toxoplasma gondii*. While previous studies have provided high-resolution views of the parasite’s invasion machinery, MyoH has remained elusive at the nanoscale. We resolved differences in radial organization between the N- and C-termini of the motor, thus determining the orientation of the protein in the apical space. Two-color imaging revealed the organization of the motor in the greater context of the parasite’s invasion complex. 3D single-molecule imaging in gel-expanded samples revealed an increase in labeling efficiency but perturbed localization of only the MyoH C-terminus, highlighting the nuanced effects of gel expansion on protein organization.

## Introduction

Apicomplexan parasites comprise a large phylum of obligate intracellular eukaryotic pathogens that includes the causative agents of malaria (*Plasmodium* spp.), cryptosporidiosis (*Cryptosporidium* spp.), and toxoplasmosis (*Toxoplasma gondii*). These parasites impose a substantial burden on human health worldwide (1–4). They also infect a wide range of domesticated and wild animals, resulting in significant veterinary and economic impact (5). *Toxoplasma gondii*, the causative agent of toxoplasmosis, has been estimated to infect about a third of the global human population (6,7). Infection is frequently asymptomatic but can cause severe disease when acquired or reactivated in immunocompromised individuals and during congenital transmission (8–10). A defining feature of apicomplexan biology is the reliance on repeated cycles of host-cell invasion, intracellular replication, and parasite egress. Invasive stages are highly polarized and deploy specialized cytoskeletal and secretory structures at the parasite apex to coordinate entry into host cells (11). *Toxoplasma gondii* belongs to the coccidian subgroup of apicomplexans, which possess a distinctive apical cytoskeletal structure known as the conoid (Fig. 1A).

**Figure 1:**
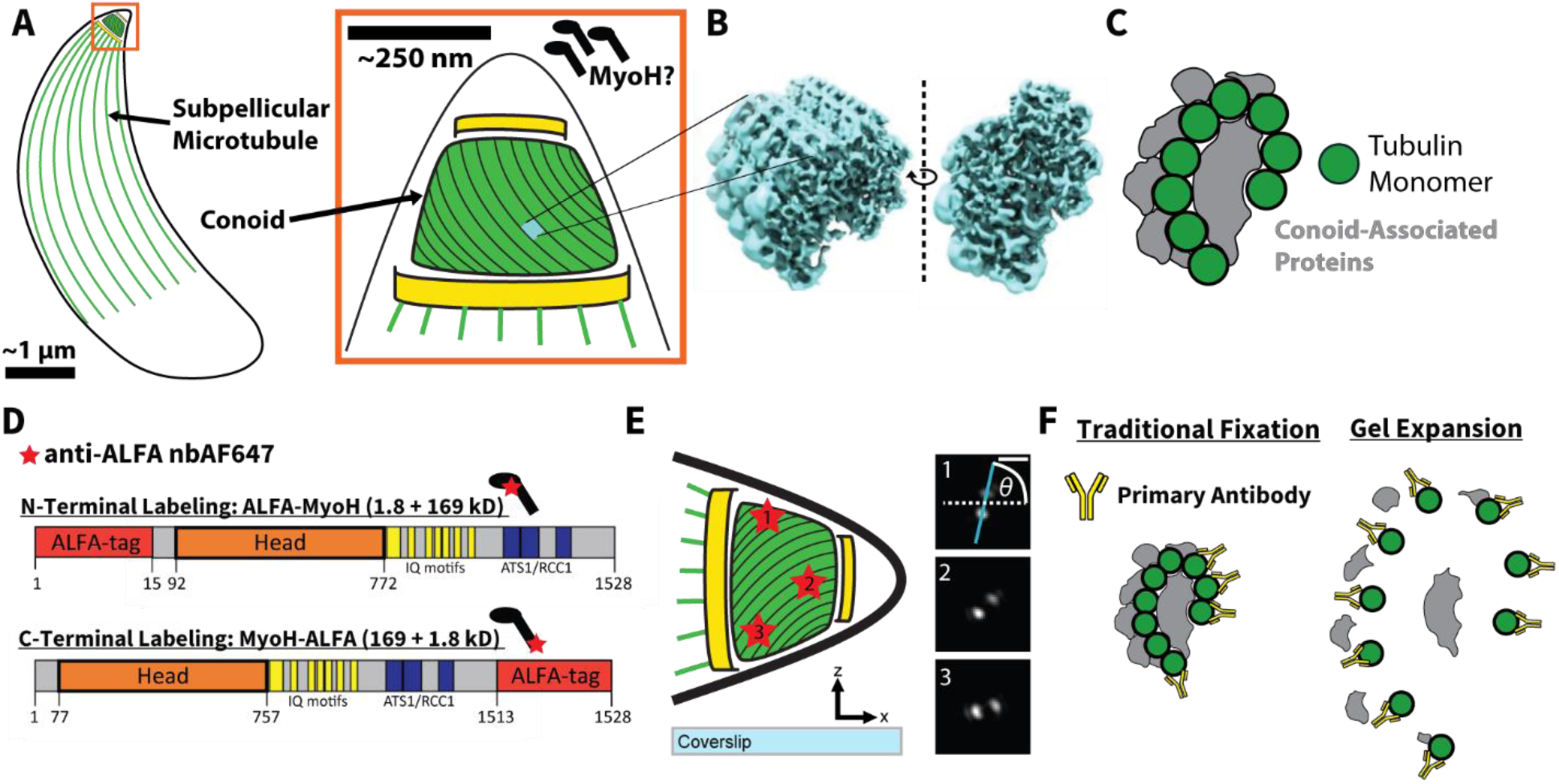
3D Single-Molecule Super-Resolution Microscopy of the Apical Complex in Traditionally Fixed and Gel-Expanded Parasites. A) Toxoplasma Gondii utilizes an apical complex to drive invasion of host cells. The conoid is a prominent feature of the apical complex. The precise organization of MyoH within the apical complex is unknown. B) Cryo-electron tomography has provided a density map of the conoid fibrils in situ (modified from Ref. (19)). C) The conoid fibrils are comprised of a comma-shaped arrangement of tubulin monomers with several conoid-associated proteins coating the tubulin. D) Two lines of parasite are created to label either terminus of MyoH, as indicated by the relative location of the red star on the miniaturized cartoon of a myosin monomer. MyoH-ALFA possesses a C-terminal fusion of the ALFA tag, while ALFA-MyoH possesses an N-terminal fusion. The ALFA tag is targeted by a nanobody conjugated to Alexa Fluor 647. E) The Double-Helix Point Spread Function (DHPSF) allows for scanning-free imaging of single-molecule fluorescence over a ∼2.5-μm axial range. Fluorophores (red stars 1, 2, and 3) have their z position encoded in the angle (θ) formed between the line bisecting the two lobes of the PSF (cyan) and the horizontal (dashed white). F) Gel expansion has been shown to increase protein accessibility, which is useful for the dense protein environment of the apical complex. Scale Bar: (E) 1 μm.

The conoid is positioned at the extreme apex of the parasite and is part of the coccidian apical complex. It is composed of unusual tubulin polymers known as conoid fibrils, which assemble into a tapered cone-shaped scaffold (12–14). Unlike canonical microtubules, these fibrils adopt a highly curved architecture. The conoid can undergo regulated protrusion and retraction during invasion-related behaviors and is associated with several additional apical structures, including the preconoidal rings, intraconoidal microtubules, and the apical polar ring (15–17). Together, these elements provide a cytoskeletal framework for organizing secretory organelles and for mechanical activity at the parasite apex (11,18). Recent cryo-electron microscopy and cryo-electron tomography studies have refined our view of conoid architecture (Fig. 1B) and revealed a complex set of associated proteins (19,20) (Fig. 1C). However, how this structure contributes to force generation remains poorly understood.

Actomyosin motors play central roles in parasite motility and invasion (21,22). In *T. gondii*, the best characterized motor is MyoA, which functions along the parasite periphery as part of the glideosome associated with the inner membrane complex (23,24). A second unconventional myosin, MyoH, localizes specifically to the conoid and is essential for gliding motility, invasion, and egress (25,26). Parasites lacking MyoH show severe defects in these processes despite retaining an intact MyoA-based glideosome, indicating that MyoH performs a distinct role at the parasite apex (25,27). MyoH contains an extended neck and tail region with multiple ATS1/RCC1 domains, and these regions are required for targeting the motor to the conoid. Current models propose that MyoH functions at the start of an apical force-transmission pathway that subsequently engages MyoA along the parasite body (25).

Despite its clear association with the conoid, the spatial organization of MyoH within this structure remains uncharacterized. The conoid is composed of multiple fibrils arranged around the parasite apex, forming a tapered scaffold that undergoes protrusion and retraction during invasion. How MyoH is positioned relative to this scaffold is not known. The orientation of the motor with respect to the conoid fibrils, its radial placement within the conoid, and its distribution along the conoid axis are all expected to influence how forces generated by MyoH are transmitted during invasion-related motility. In addition, it is not known whether MyoH interacts directly with tubulin polymers or is recruited through other conoid-associated proteins.

Resolving the nanoscale organization of proteins within the conoid is experimentally challenging. The dimensions of the structure lie below the diffraction limit of conventional visible light microscopy, and the dense molecular environment of the parasite apex can limit epitope accessibility. To examine how MyoH is arranged relative to the conoid scaffold, we combined super-resolved, extended axial range 3D, single-molecule localization microscopy (SMLM) with gel-based expansion approaches. Specifically, we created two separate parasite strains with fusions of the ALFA tag to the N- and C-terminus of MyoH (Fig. 1D). We used the double-helix point spread function (DHPSF) and (d)STORM-style blinking (28,29) to localize single MyoH molecules in 3D with high-precision (AF647: 12 nm lateral, 20 nm axial; CF583R: 27 nm lateral, 40 nm axial; Fig. S4) across a 2.5-µm axial range (30,31). The DHPSF makes single-molecule emission appear as two spots on the camera, with the rotation angle of the lobes encoding the z position of the emitter and the midpoint between the lobes encoding the x-y position (Fig. 1E). Single-color reconstructions of either end of the protein allowed us to measure the organization and orientation of the motor in the apical space. Two-color 3D imaging with tubulin localized MyoH relative to the conoid. We found conoid labeling to be sparse in traditionally fixed samples and thus turned to gel expansion, which has been combined with SMLM not only for improved resolution (32,33) but also for decreasing molecular crowding and increasing epitope accessibility (34) (Fig. 1F). 3D single-molecule reconstructions of MyoH and tubulin in gel-expanded samples revealed improved tubulin labeling but also suggested MyoH organization is perturbed by gel expansion. These experiments allowed us to determine the spatial distribution of MyoH at the parasite apex and to evaluate how labeling constraints influence the interpretation of nanoscale organization within the conoid.

## Materials and Methods

### Parasite Maintenance and Cell Culture

Toxoplasma gondii RHΔhxgprt and RHΔ*hxgprt*Δku80 strains were maintained by growth in confluent human foreskin fibroblasts (HFFs; ATCC, CCD-1112Sk) in Dulbecco’s modified Eagle’s medium (DMEM; Gibco, 11960069) supplemented with 10% fetal bovine serum (FBS; Corning, 35-016-CV), 2 mM glutamine (Corning, 5-005-CI), and 100 U/mL penicillin: 100 μg/mL streptomycin (MP Biomedicals, 091670249) at 37°C in 5% CO2.

### Plasmids and Parasite Line Generation

See Table S1 for all primers and oligonucleotides used and Table S2 for all plasmids. All sgRNAs were cloned into the pSAG1:U6-Cas9:sgUPRT (35). pBAD/HisB-PAmKate was a gift from Vladislav Verkhusha (Addgene plasmid 3269). All cloning steps were performed using restriction-free (RF) cloning by generating megaprimers containing homology regions to the target plasmid sequence.

PAmKate (36) was PCR-amplified from pXyl_PAmKate_PopZ (37) using primers 20 and 21 and cloned into pSAG1:U6-Cas9:sgUPRT. PAmKate together with the SAG1 3′ UTR was subsequently amplified using primers 24 and 25 and cloned into pGRA1-HXGPRT-3xHA. A linker sequence (ggaggaggaagt)_2_ was introduced upstream of PAmKate using primers 64/65, and a stop codon was added downstream using primers 42/43, generating plasmid pGRA1-PAmKate-HXGPRT.

To generate a construct for endogenous C-terminal tagging of *MyoH* (TGGT1_243250), genomic fragments corresponding to the MyoH C-terminus and MyoH 3′ UTR were PCR-amplified using primers 66/67 and 68/69, respectively, and cloned into pGRA1-PAmKate-HXGPRT, generating pGRA1-MyoH-PAmKate-HXGPRT. PAmKate was subsequently replaced with an ALFA tag using primers 169 and 170, generating pGRA1-MyoH-ALFA-HXGPRT.

For parasite transfection, the tagging cassette containing the homology regions, linker–PAmKate or ALFA tag, and the selection marker was PCR-amplified using primers 127 and 234. A total of 10 µg of this PCR product together with 10 µg of pSAG1:U6-Cas9:sgMyoH 3′UTR were transfected into RHΔ*hxgprt*Δ*ku80* parasites.

For N-terminal tagging, plasmid pGRA1-HXGPRT-R-ALFA-MyoH was generated using the same cloning strategy described above in the reduced backbone pGRA1-HXGPRT-R (derived from pGRA1-HXGPRT-3xHA) and transfected together with pSAG1:U6-Cas9:sgMyoH_5′UTR into RHΔ*hxgprt* parasites.

### Parasite Preparation for Imaging

HFF monolayer infected with Toxoplasma tachyzoites was washed twice with PBS and once with phenol red-free DMEM (Gibco, 31053028) supplemented with 2 mM glutamine, 100 U/mL penicillin and 100 μg/mL streptomycin. The tachyzoites were released from the HFFs by mechanically disrupting the monolayers with disposable scrapers and passing the material through a 25-gauge syringe. The parasites were added to fresh monolayers of HFFs (which were pre-washed with PBS and phenol red-free DMEM). After 18-20 hours, the infected cells were washed twice with PBS and once with Endo buffer (EB), containing 45 mM potassium sulfate, 106 mM sucrose, 10 mM magnesium sulfate, 20 mM Tris buffer at pH 7.2, 5 mM glucose, and 0.35% bovine serum albumin (38). The HFFs monolayers were scraped and passed through a 25-gauge syringe, then a 27-gauge syringe, and finally filtered with a 5 μm syringe filter (Millex, SLSV025LS).

To stimulate conoid protrusion, HFFs were washed with Hank’s balanced salt solution (HBSS; Gibco, 14175103), supplemented with 1 mM MgCL_2_, 1 mM CaCl_2_, 10 mM NaHCO_3_, and 20 mM HEPES, pH 7. The HFFs monolayers were scraped and passed through a 25-gauge syringe, then a 27-gauge syringe, and finally filtered with a 5 μm syringe filter. Calcium-ionophore (Sigma, C7522) was added to the sample at a final concentration of 1 μM and then incubated for 10 minutes at room temperature.

Tachyzoites were placed into imaging chambers (Cellvis, D29-20-1.5H) precoated with poly-L-lysine and centrifuged to facilitate attachment to the glass bottom. Samples were first prefixed and permeabilized for 2 minutes using 0.3% (v/v) glutaraldehyde and 0.25% (v/v) Triton X-100 in cytoskeleton buffer (10 μM MES pH 6.1, 150 mM NaCl, 5 mM EGTA, 5 mM D-glucose, 5 mM MgCl). Samples were fixed for 10 minutes using 2% (v/v) glutaraldehyde in cytoskeleton buffer, then quenched for 7 minutes with 0.1% (w/v) NaBH4 in PBS.

For MyoH-PAmKate imaging, samples were fixed for 15 minutes using 3% (v/v) paraformaldehyde in PBS, then quenched for 7 minutes with 0.1% (w/v) NaBH4 in PBS.

### Fixed Cell Labeling

After fixation, single-color MyoH samples were labeled with FluoTag-X2 anti-ALFA nanobodies conjugated to Alexa Fluor 647 (NanoTag Biotechnologies, N1502-AF647-L) using a 1:500 dilution in 2% BSA (Fisher Scientific, PI37525) in PBS for 2 hours at room temperature. Samples were then washed 3 times for 5 minutes each time with 2% BSA (Fisher Scientific, PI37525) in PBS. Samples were then incubated with a 1:50,000 dilution of 200 nm 580/605 nm FluoSpheres (Thermo Fisher Scientific, F8810) in PBS for 15 minutes and then rinsed with PBS 3 times for 5 minutes each time.

Two-color fixed cell samples were first incubated with mouse monoclonal anti-α tubulin (Thermo Fisher, 32-2500) and mouse monoclonal anti-β Millipore Sigma, T5293) both diluted to a concentration of 2 µg/ml in 2% BSA in PBS overnight at 4 °C. Samples were then washed 3 times with 5-minute incubations with 2% BSA in PBS. Washing was then followed by the single-color fixed cell labeling procedure detailed above with the following changes. CF583R goat anti-mouse secondary antibodies (Biotium, 20903-500µl) were added to the nanobody solution at a dilution of 1:500. A 1:12.5 dilution of 100 nm gold nanoparticles (Cytodiagnostics, G-100-20) in PBS was substituted for FluoSpheres as fiducials.

Just before imaging, labeled fixed cell samples were filled with freshly prepared blinking buffer and sealed to eliminate contact of the buffer with an air interface. The blinking buffer consisted of 200 U/ml glucose oxidase, 1000 U/ml catalase, 10% w/v glucose, 200 mM Tris-HCl pH 8.0, 10 mM NaCl. (See Supplemental Methods for further details).

### Gel Expansion

Fixed parasite samples were gel expanded using an adaptation of a previously published protocol (39). Full details can be found in Supplemental Methods. Each parasite sample was incubated in a humid chamber with 1 ml of FA solution (0.7% formaldehyde, 1% acrylamide in PBS) for 5 hours at 37 °C. Samples then had FA solution removed, were gently blown dry with nitrogen, and then placed in humid chambers on ice. 35 μl of monomer solution (19% sodium acrylate, 10% acrylamide, 0.1% BIS, 0.5% ammonium persulfate, 0.5% TEMED) was placed onto each parasite sample, and a 12-mm diameter circular coverslip (Fisher Scientific, 12-541-001) was placed on top of each drop of monomer solution. The humid chamber was then closed and incubated on ice for 5 minutes followed by incubation at 37 °C for 1 hour. Gels were then incubated with denaturation buffer (200 mM SDS, 200 mM NaCl, 50 mM Tris-Base, pH 9) on an orbital shaker with gentle agitation for 30 minutes followed by incubation in denaturation buffer at 95 °C for 1.5 hours. Each gel was then incubated twice for 30 minutes with 150-200 ml of pure water and left overnight in pure water.

### Gel Labeling and Sample Chamber Preparation

After overnight incubation in water, gels were incubated twice for 30 minutes in fresh PBS. Gels were then cut to size to fit on coverslips. Single-color MyoH samples were labeled with FluoTag-X2 anti-ALFA nanobodies conjugated to Alexa Fluor 647 (NanoTag Biotechnologies, N1502-AF647-L) using a 1:500 dilution in 2% BSA (Fisher Scientific, PI37525) in PBS for 2 hours at 37 °C. Samples were then washed 3 times with 10-minute incubations with 0.1% Tween-20 (Millipore Sigma, P9416) in PBS. An orbital shaker was used during all gel washes to gently agitate the gels in washing solution.

Two-color gel samples were also incubated twice for 30 minutes in fresh PBS and cut to fit on coverslips. Gels were then incubated with mouse anti-α tubulin primary antibodies (ABCD Antibodies, ABCD_AA345) and mouse anti-β tubulin primary antibodies (ABCD Antibodies, ABCD_AA344) both diluted to a concentration of 2 µg/ml in 2% BSA in PBS for 3 hours at 37 °C. Samples were then washed with 3 times with 10-minute incubations with 0.1% Tween-20 in PBS with gentle agitation. Washing was then followed by the single-color gel labeling procedure detailed above with the addition of CF583R goat anti-mouse secondary antibodies (Biotium, 20903-500µl) to the nanobody solution at a dilution of 1:500.

A gel piece was then selected for imaging and placed onto the coverslip of a clean CellVis chamber. The gel was then covered with freshly prepared blinking buffer and sealed with a larger circular coverslip as described above. After 1 hour, the gel was unsealed and the blinking buffer removed from the chamber. The gel was then transferred to a poly-L-lysine and fiducial bead-treated CellVis chamber (See Supplemental Methods). Three small, circular coverslips (Fisher Scientific, 12-541-001) that were glued together (Gorilla Clear Grip) were placed on top of the gel piece to hold the gel piece onto the coverslip. Fresh blinking buffer was added to the chamber, and the chamber was then sealed with a larger coverslip as described in Supplemental Methods.

### 3D DHPSF Imaging

3D DHPSF (d)STORM imaging was performed with a previously described, homebuilt, two-color DHSPF microscope (40,41) controlled using the PYthon Microscopy Environment (PYME) (42) (Fig. S3). See Supplemental Methods for full microscope details.

Single-color GA-fixed and gel-expanded MyoH imaging was performed by first shelving the MyoH signal with 33 W/cm^2^ intensity 647 nm illumination for 1-2 minutes until isolated DHPSFs were visible followed by single-molecule blinking and imaging with 3.3 kW/cm^2^ 647 nm illumination and increasing 405 nm illumination (see Supplemental Methods) until fully bleached (∼23,000 frames) with camera parameters of 39 ms exposure and 284X EM gain.

Two-color imaging of MyoH and tubulin was performed sequentially with MyoH imaging occurring first followed by tubulin imaging. MyoH imaging in this case was performed similarly to single-color MyoH imaging with the following changes. Imaging was limited to 10,000 frames and with minimal 405 nm photoactivation (see Supplemental Methods) to limit photobleaching of the tubulin signal. Tubulin was then imaged by first shelving with 66 W/cm^2^ intensity 561 nm illumination until isolated DHPSFs were visible followed by single-molecule imaging with 6.6 kW/cm^2^ 561 nm illumination and increasing 405 nm illumination (see Supplemental Methods) until fully bleached with camera parameters of 15 ms exposures and 284X EM gain. The same two-color imaging protocol was used for both GA-fixed and gel-expanded samples.

After cellular imaging, a calibration sample of 100 nm TetraSpeck beads (Invitrogen, T7279) in 1% agarose was used to acquire 3D bead localizations simultaneously in both color channels to compute a registration between channels. Finally, a bead was scanned axially in 50-nm steps across a z range of 3.2 μm to generate a θ-to-z calibration.

### Localizing and Processing DHPSF Data

DHPSF images were localized using a Python-based plugin for PYME (43). Localizations were then assigned 𝑧 positions using a spline of the z stack calibrations, filtered on localization precision and DHPSF parameters, registered using a locally-weighted mean quadratic transformation (for two-color data) (44), drift corrected using smoothed fiducial traces, foreshortening corrected with a linear factor of 0.7, and merged. See Supplemental Methods and Ref. (45) for full details and processing scripts for localization, filtering, merging, and registration. Median localization precisions by dye and sample preparation were as follows: AF647, GA-fixed: 12 nm 𝑥, 20 nm 𝑧; AF647, Gel-Expanded: 19 nm 𝑥, 30 nm 𝑧; CF583R GA-fixed: 27 nm 𝑥, 40 nm 𝑧; CF583R Gel-Expanded: 25 nm 𝑥, 37 nm 𝑧 (Fig. S4). Localizations were rendered as pointsprites in PYMEVisualize (46).

### Single-Color Apical Localization Processing and Quantifications

Apical localizations were then isolated and aligned. For each field of view, the 𝑥, 𝑦, and 𝑧 centers of apical localizations were manually selected. Localizations within a specified cropping radius from each center were then isolated, with a cropping radius of 250 nm and 500 nm for GA-fixed and gel-expanded localizations, respectively (before expansion factor correction for gel localizations). Isolated apical localizations were then aligned to the same coordinate system using principal component analysis (PCA) with the 𝑧 axis representing the cell axis. Localizations were then filtered to eliminate the bottom and top 1.5% along the cell axis for each set of apical localizations. In addition to real space 𝑧 position, apical localizations were assigned normalized 𝑧 coordinates computed with the following formula:

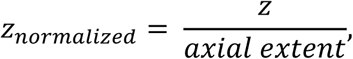

A sliding window along the normalized 𝑧 coordinate (width: 0.1, increment: 0.01) was used to quantify radius as a function of the cell axis. For each window, the median of localizations within that window was assigned as the radius. A sliding window along the normalized 𝑧 coordinate (width: 0.05, increment: 0.005) was also employed to quantify localization density along the cell axis. For each window, the localization density was computed as

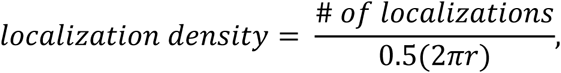

where 𝑟 is the median radius of the bin, to account for the surface area represented by each window. See Ref. (45) for apical localization processing scripts. 95% confidence intervals were calculated using the MATLAB function *ttest2* (MATLAB version 2025a).

### Two-Color Apical Localization Processing and Quantifications

Two-color localizations were processed similarly to single-color localizations with the following modifications. The cropping radius was 300 nm and 500 nm for GA-fixed and gel-expanded localizations, respectively (before expansion factor correction for gel localizations). Localizations from both color channels were considered for 𝑚𝑎𝑥(𝑧) and 𝑚𝑖𝑛(𝑧) for cell axis normalization. Two-color reconstructions were further filtered radially with 50 nm < radius < 250 nm GA-fixed data and 120 nm < radius < 550 nm for gel-expanded data (before expansion factor correction) after PCA alignment. Sliding-window radial quantification of localizations was modified such that for each MyoH localization, the corresponding tubulin radius was computed as the median radius of tubulin localizations within a window of 𝑧 = 0.2 along the normalized cell axis and within an angular window of 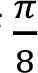 𝑟𝑎𝑑 in the 𝑥-𝑦 plane. See Ref. (45) for apical localization processing scripts.

### Gel Expansion Factor Correction

To compare GA-fixed apical reconstructions to those from gel-expanded samples, we first computed an expansion factor using the tubulin localizations. While it is common to simply measure the change in the size of the gel to compute an expansion factor, it is documented that the measured size of biological structures in gel-expanded samples often deviates from the size expected from macroscopic gel expansion measurements (47). Cryo-electron microscopy of unfixed parasites has been used to measure the base of the conoid to have a radius of 190 nm (Fig. S5D). The conoid base was thus chosen as an internal reference to determine expansion factor for each gel. We isolated tubulin localizations from 0 to 0.2 on the normalized cell axis and divided the median radius of these localizations by 190 nm to approximate the expansion factor. All 𝑥, 𝑦, 𝑧, and radial coordinates for gel-expanded apical reconstructions were then divided by the expansion factor.

## Results

### Single-Color MyoH Reconstructions Resolve Outward Orientation of the N-Terminus

To visualize MyoH within parasites, we created two separate lines containing endogenously expressed MyoH fused to the ALFA tag, one lines with an N-terminal fusion (ALFA-MyoH) and the other with a C-terminal fusion (MyoH-ALFA) (Fig. 1D, Fig. S1A-B). The ALFA tag is a 15 amino acid sequence forming an alpha helix that can be targeted with high specificity by a small, dye-conjugated nanobody (48). Given that MyoH is indispensable for host cell invasion (25), the ALFA tag is an appealing choice due to its small size (∼2 nm), which minimizes perturbation to the protein. Additionally, the use of primary and secondary antibodies increases the measured, effective size of labeled structures due to the size of the antibodies themselves (49–51); our use of nanobodies here drastically reduces this distortionary effect (51).

After confirming specific labeling of MyoH (and viability) in both lines (Fig. S2A-C), we performed 3D (d)STORM imaging of glutaraldehyde (GA)-fixed, conoid-protruded and conoid-retracted parasites with a homebuilt DHPSF microscope (Fig. S3A). The microscope uses a phase mask to modulate the microscope response such that point sources appear on the detector as two Gaussian lobes. The midpoint between the lobes provides an estimate of emitter 𝑥-𝑦 position, while the angle between the reference axis and the line connecting the lobes encodes the 𝑧 position (52) (Fig. 1E and Fig. S3B, C). The DHPSF allows for the localization of single-molecule fluorescence in 3D over an extended axial range (2.5 µm) without the need for axial scanning, proving especially useful for imaging many parasites in a field of view, each with a varied orientation on the coverslip.

Imaging conoid-protruded MyoH-ALFA parasites with the DHPSF revealed a dense set of localizations with precisions beyond the diffraction limit from the apical end of the parasite as expected (Fig. 2A). Rotating these localizations in 3D, we see that the localizations form a truncated cone when viewed orthogonally to the cell axis and a ring-shaped profile when viewed down the cell axis (Fig. 2B). This localization pattern was also observed for ALFA-MyoH (Fig. S5A, B). As a result of the consistent shape of the apical localizations in both parasite lines, we were able to align the apical localizations to a common set of cellular coordinates (Fig. 2C, D).

**Figure 2:**
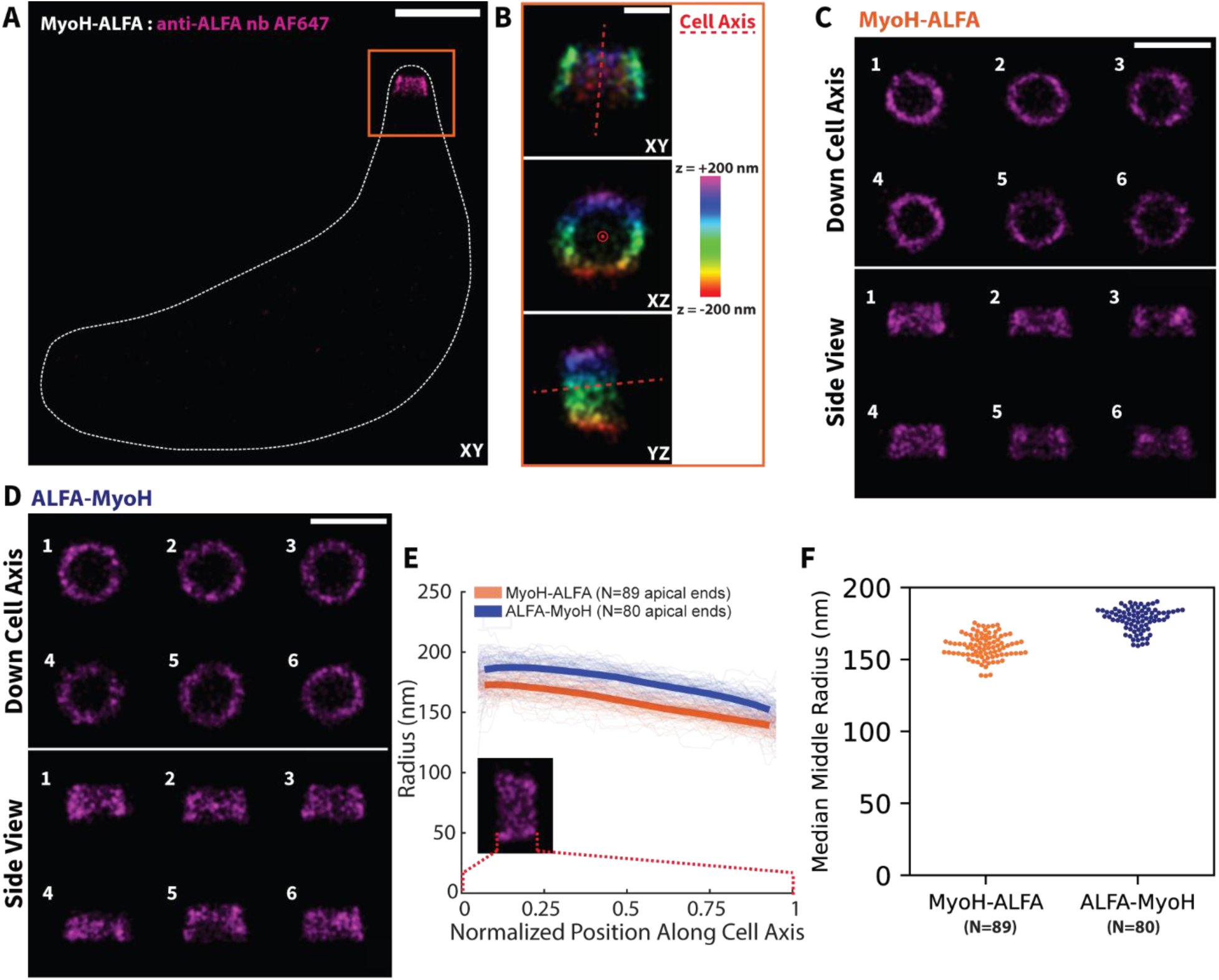
3D Reconstructions of Apical MyoH-ALFA and ALFA-MyoH Reveal Differential Organization of the MyoH N- and C-Termini: A) XY projection of localizations from a representative conoid-protruded MyoH-ALFA parasite with the parasite membrane marked with white, dashed line. A dense set of localizations is seen at the apical tip of the parasite. B) Apical MyoH-ALFA localizations for a representative conoid-protruded parasite viewed in XY, XZ, and YZ projections and colored by z. C) Six conoid-protruded apical MyoH-ALFA localizations (labeled 1-6) that have been PCA-aligned to a common coordinate system and viewed from two different orientations: down the cell axis (top two rows) and a side view (bottom two rows) D) As for (C) but for conoid-protruded apical ALFA-MyoH localizations. E) Radius vs normalized position along cell axis for conoid-protruded MyoH-ALFA (orange) (N=89 cells) and ALFA-MyoH (blue) (N=80 cells) localizations. Light lines represent binned radial values for single cells. Dark lines represent the median radius for all cells with the linewidth representing the standard error of the mean. F) Median middle radius of each apical end for MyoH-ALFA (orange) (N=89 cells) and ALFA-MyoH (blue) (N=80 cells) from conoid-protruded parasites. Each dot shows the median radius of localizations ranging from 0.45 to 0.55 along the normalized cell axis for a single cell. Scale Bars: (A) 1 μm, (B) 200 nm, (C-D) 500 nm.

After alignment, we quantified various aspects of the apical MyoH structures to determine which parameters if any differed between the ALFA-MyoH and MyoH-ALFA localizations. We found the axial extent, or height along the cell axis, of ALFA-MyoH reconstructions to be larger than that of MyoH-ALFA reconstructions. In conoid-protruded parasites, ALFA-MyoH and MyoH-ALFA localizations had median axial extents of 253 nm and 229 nm, respectively (95% C.I. difference of 27 ± 6 nm) (Fig. S5C). In conoid-retracted parasites, ALFA-MyoH and MyoH-ALFA localizations had median axial extents of 247 nm and 218 nm (95% C.I. difference of 29 ± 6 nm) (Fig. S5D). (Interestingly, conoid-protruded localizations had slightly larger axial extents relative to conoid-retracted localizations (MyoH-ALFA: 9 ± 5 nm; ALFA-MyoH: 6 ± 6 nm; 95% C.I. difference).) These results suggest the motor head spans a greater range along the cell axis relative to the motor tail.

We then quantified the radius of MyoH localizations relative to the cell axis. Pooling localizations from all apical ends, ALFA-MyoH localizations peaked at a larger radius than MyoH-ALFA localizations for both conoid-protruded and conoid-retracted parasites (Fig. S5E, F). To better understand this difference, we measured the MyoH radius as a function of position along the cell axis (Fig. 2E). Since apical reconstructions from each parasite line differed in height, position along the cell axis was normalized such that 1 represented the apical tip of the localizations and 0 the opposite end. Both the ALFA-MyoH localizations and MyoH-ALFA localizations show decreasing radii moving from 0 to 1 along the normalized cell axis, as is expected from the visible tapering of the structures (Fig. 2E). However, ALFA-MyoH localizations consistently have a larger radius than MyoH-ALFA localizations along the entirety of the cell axis in both conoid-protruded (Fig. 2E) and conoid-retracted parasites (Fig. S5G).

To quantify this difference, we measured each cell’s median middle MyoH radius as the median radius of localizations ranging from 0.45 to 0.55 along the normalized cell axis. This central window accounts for differences in axial extent and the tapering of the structure. In conoid-protruded parasites, MyoH-ALFA cells had a middle radius of 159 ± 8 nm (mean ± s.d.), while ALFA-MyoH cells had a middle radius of 178 ± 8 nm (mean ± s.d.), representing a difference in middle radius of 19 ± 3 nm (95% C.I.) (Fig. 2F). Similarly, in conoid-retracted parasites, MyoH-ALFA cells had a middle radius of 160 ± 7 nm (mean ± s.d.), while ALFA-MyoH cells had a middle radius of 174 ± 9 nm (mean ± s.d.), representing a difference in middle radius of 14 ± 3 nm (95% C.I.) (Fig. S5H). Thus, on average, the N-terminus of MyoH lies radially outward of the C-terminus in the apical complex.

### Two-Color Imaging Determines MyoH Position Relative to the Conoid

While single-color MyoH imaging has revealed the orientation of the protein’s termini in the apical complex, two-color imaging is necessary to localize the motor within the greater context of the conoid. The tubulin-based conoid represents a prominent and well-characterized structure in the apical complex (19,20), making tubulin localizations in the conoid an appealing landmark to provide context for MyoH localizations. For the remainder of this work, all data are from conoid-protruded parasites only.

GA-fixed MyoH-ALFA and ALFA-MyoH parasites were labeled for both tubulin and MyoH and imaged with the DHPSF. We found tubulin labeling in the conoid to be highly heterogenous (Fig. 3A-C), even when labeling with a variety of different anti-tubulin antibodies (see Supplemental Methods). In some cases, apical tubulin labeling heavily favored the conoid base (Fig. S6B), while in others a more faithful reconstruction of the conoid was recovered (Fig. S6C). This labeling bias for the conoid base is clearly visible when localization density is plotted along the normalized cell axis (Fig. 3D, E). For each apical end, the number of localizations in each color channel was binned along the normalized cell axis and subsequently normalized to account for the surface area represented by each bin. In both MyoH-ALFA and ALFA-MyoH parasites, tubulin localizations were denser at the base of the conoid and gradually decreased along the cell axis. MyoH localizations in both parasite lines showed a different pattern with localization density increasing from the base of the conoid to the tip of the conoid. Given the dense protein composition of the conoid (19,20), it is likely that steric hindrance limits the accessibility of anti-tubulin antibodies, thus leading to poorly sampled conoid reconstructions. As a result, when looking at 50-nm cross sections orthogonal to the cell axis (Fig. 3B, C, bottom rows), we see that some cross sections show more tubulin localization than others to serve as a reference point for MyoH localizations.

**Figure 3:**
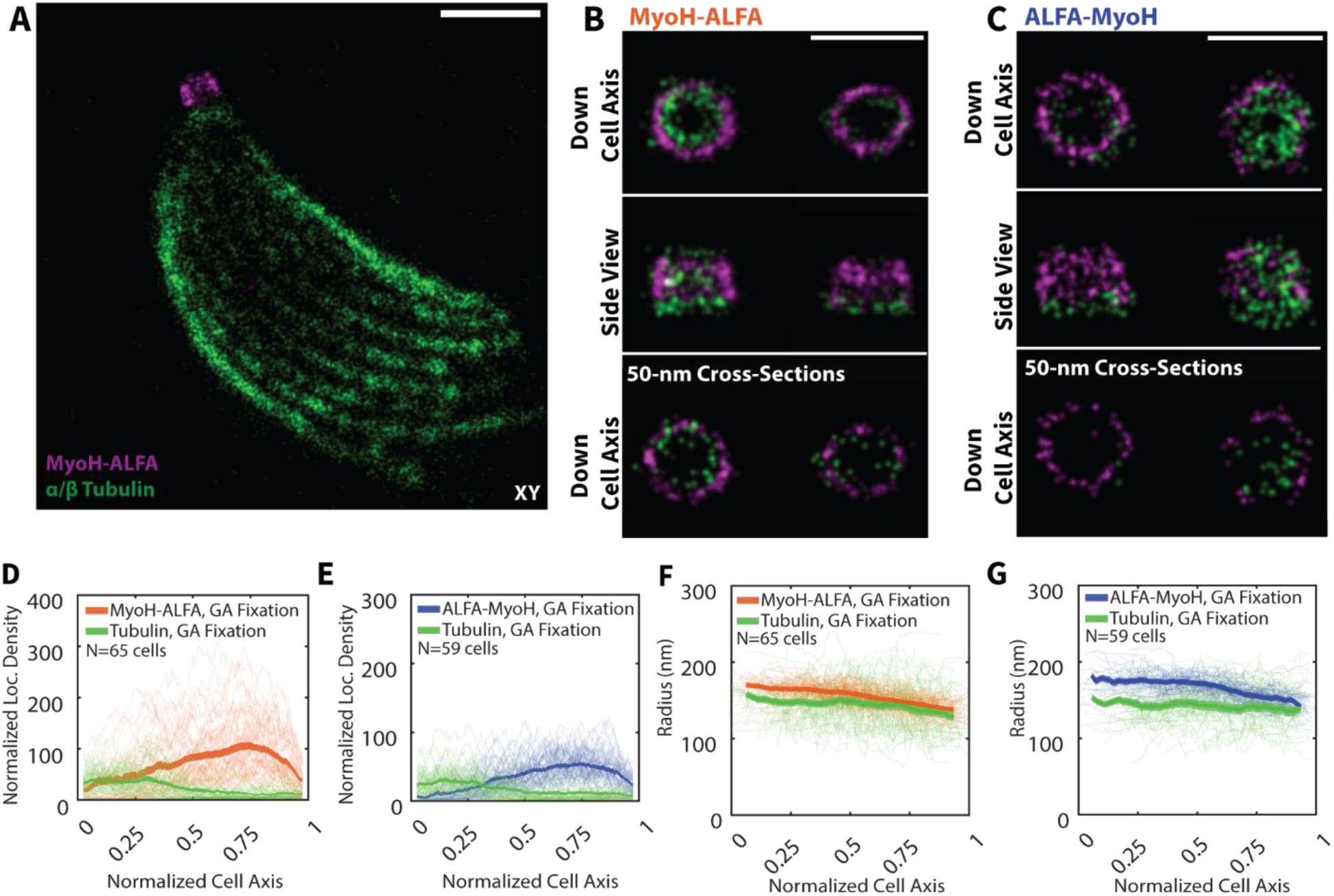
Two-Color Reconstructions in GA Fixed Samples Reveal Organization of MyoH Relative to the Conoid Fibrils. A) XY projection of two-color, 3D reconstruction of tubulin (green) and MyoH-ALFA (magenta) in a GA fixed MyoH-ALFA parasite. B) Two aligned, two-color apical reconstructions of MyoH-ALFA (magenta) and tubulin (green) shown in two orientations: down the cell axis (top row) and side view (middle row). Bottom row shows a cross-section 50-nm thick along the cell axis and viewed down the cell axis. C) Two aligned, two-color apical reconstructions of ALFA-MyoH (magenta) and tubulin (green) shown in two orientations: down the cell axis (top row) and side view (middle row). Bottom row shows a cross-section 50-nm thick along the cell axis and viewed down the cell axis. D-E) Normalized localization density vs normalized cell axis for tubulin (green) and MyoH-ALFA (D; N=65 cells) (orange) or ALFA-MyoH (E; N=59 cells) (blue). Light lines represent localization density from single cells. Dark lines show the median localization density for all cells with the linewidth representing the standard error of the mean. F-G) As for (D-E), except median, binned radius vs position along the cell axis for tubulin (green) and MyoH-ALFA (F; N=65 cells) (orange) or ALFA-MyoH (G; N=59 cells) (blue). Scale Bars: (A) 1 μm, (B-C) 500 nm.

To mitigate the effects of uneven conoid labeling, the radius of each MyoH localization was compared to that of tubulin localizations in close proximity both along the cell axis and angularly in the plane orthogonal to the cell axis. We see that MyoH-ALFA localizations are located slightly radially exterior of the tubulin localizations (13 nm median radial difference), with ALFA-MyoH localizations even further radially relative to tubulin (25 nm median radial difference) (Fig. 3F-G). The radial difference between the C-terminus and tubulin is within the ∼18-nm offset distance expected for labeling with primary and secondary antibodies (51). Thus, we cannot predict how the C-terminus is positioned relative to the tubulin. The median radius of all conoid localizations <0.1 along the normalized cell axis, representing the base of the conoid, was 157 nm. The expected base conoid radius is approximately 190 nm measured by cryo-electron microscopy (Fig. S6D).

The relatively small radius of conoid localizations suggests preferential labeling of tubulin monomers of the conoid’s interior face. The reconstructions do not have the resolution, however, to determine which specific monomers are successfully labeled. Additionally, the direction in which the primary and secondary antibodies are extending from the tubulin monomers is presumably random which also limits the precision with which the dimensions of the tubulin-based components can be inferred. As a result, there is a range of possible MyoH positions relative to the tubulin filaments that are consistent with the observed radial arrangement of proteins (Fig. 6E).

### Gel-Expansion Improves MyoH Epitope Accessibility

While imaging of GA-fixed parasites revealed details of MyoH organization, the known molecular crowding in the apical complex leads to the possibility that only a fraction of the MyoH in the apical complex is accessible for labeling under GA fixation, especially considering the limited tubulin accessibility in the conoid. Gel-based sample expansion has been shown to decrease molecular crowding and thus increase epitope accessibility (34), and, when paired with SMLM, improves the effective resolution and decreases the distortionary impact of primary and secondary antibodies (32,33). Thus, we prepared GA-fixed MyoH-ALFA and ALFA-MyoH parasites and subsequently embedded them in acrylamide gels. The expanded gels were then labeled for MyoH, placed in blinking buffer, and imaged similarly to the GA-fixed MyoH samples. In the blinking buffer, gels expanded by a factor of ∼1.7X.

In both MyoH-ALFA and ALFA-MyoH parasites, we observed dense apical MyoH localizations that qualitatively appeared to take the same shape as the GA-fixed MyoH reconstructions (Fig. 4A, B). To compare GA-fixed MyoH reconstructions and gel-expanded MyoH reconstructions, radial coordinates for each cell’s apical localizations were normalized by dividing by the median radius for that cell. Comparing this normalized radius along the cell axis, we see that both the MyoH-ALFA and ALFA-MyoH localizations exhibit similar tapering, suggesting that the MyoH structure is similar in both traditional fixation and gel expansion (Fig. 4C, D). However, gel reconstructions contained ∼38% more localizations than GA-fixed localizations for MyoH-ALFA parasites and ∼88% more for ALFA-MyoH parasites (Fig. S7A), with increases in localization density at all points along the cell axis (Fig. 4E, F). These results suggest that gel expansion has increased the accessibility of both the N- and C-termini of MyoH in the apical complex.

**Figure 4:**
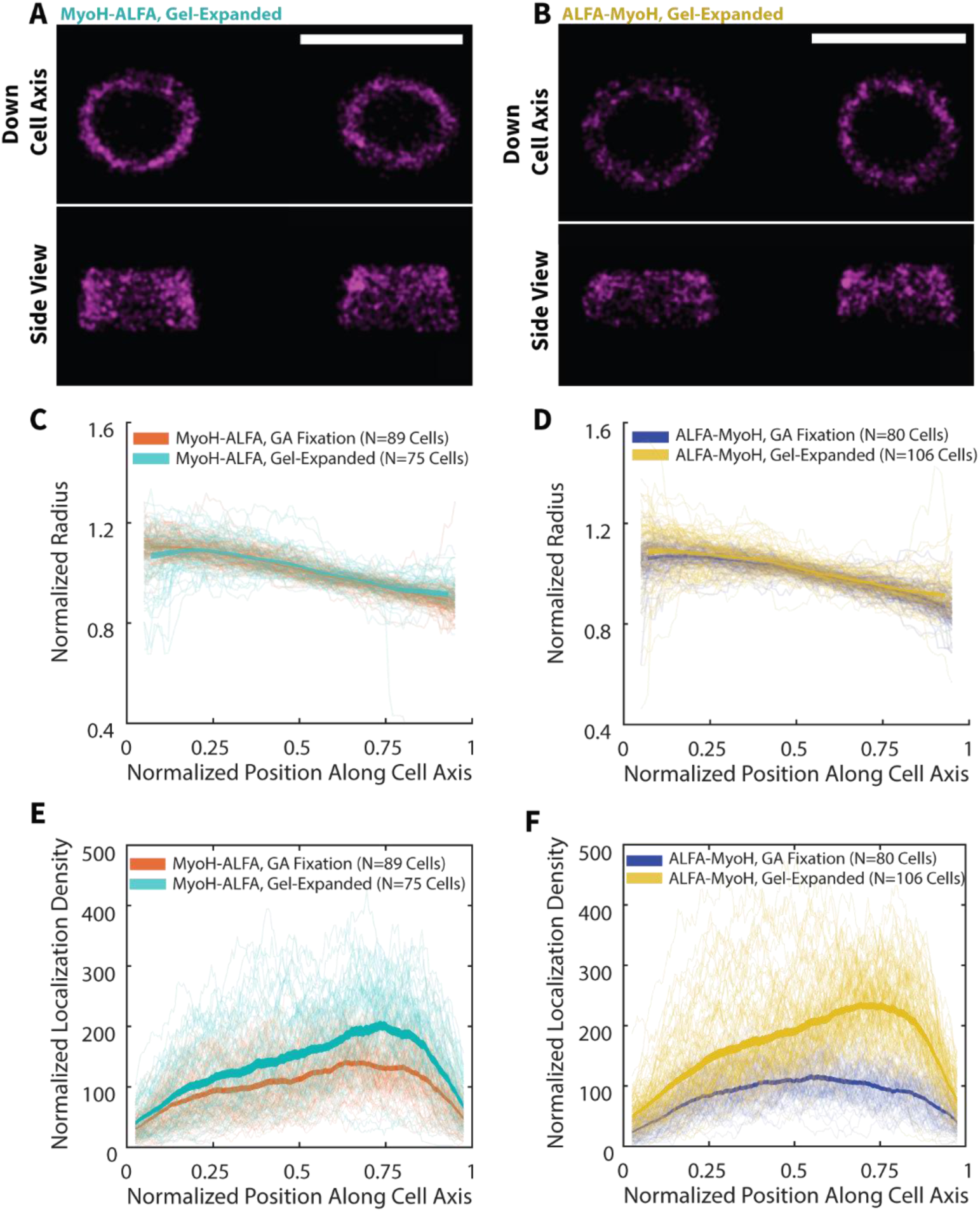
Gel Expansion Improves Accessibility of MyoH. A) Two aligned sets of apical MyoH-ALFA localizations from gel-expanded samples viewed from two different orientations: down the cell axis (top row) and side view (bottom row). B) Two aligned sets of apical ALFA-MyoH localizations from gel-expanded samples viewed from two different orientations: down the cell axis (top row) and side view (bottom row). C) Normalized radius vs normalized position along cell axis for gel-expanded MyoH-ALFA (N=75 cells) (cyan) and GA fixed MyoH-ALFA (N=89 cells) (orange) localizations. Light lines represent binned radial values for single cells. Dark lines represent the median radius for all cells with the linewidth representing the standard error of the mean. D) As for (C), except normalized radius vs normalized position along cell axis for gel-expanded ALFA-MyoH (N=106 cells) (yellow) and GA fixed ALFA-MyoH (N=80 cells) (blue) localizations. E) As for (C), except normalized localization density vs normalized position along cell axis for gel-expanded MyoH-ALFA (N=75 cells) (cyan) and GA fixed MyoH-ALFA (N=89 cells) (orange) localizations. F) As for (C), normalized localization density vs normalized position along cell axis for gel-expanded ALFA-MyoH (N=106 cells) (yellow) and GA fixed ALFA-MyoH (N=80 cells) (blue) localizations. Scale Bars: (A-B) 1 μm.

This raises the question of whether there remains a substantial fraction of MyoH that is still inaccessible to the anti-ALFA nanobody and requires expansion beyond the factor of ∼1.7X performed here. To answer this, we turned to a line of parasite with the photoactivatable, fluorescent protein PAmKate fused to the C-terminus of MyoH and endogenously expressed. This parasite line allows for labeling of essentially all MyoH in the parasite as opposed to the ALFA tag fusions which have a labeling efficiency clearly impacted by accessibility. A single-molecule fluorescence bleaching assay produced an estimate of 553 ± 90 MyoH-PAmKate molecules per apical end (mean ± s.d.) (Fig. S7B).

We can roughly estimate the number of MyoH molecules in each apical ALFA tag reconstruction to compare to the MyoH-PAmKate estimate. Quantifying the true number of molecules from a single-molecule reconstruction is difficult due to the stochastic, photophysical properties of the dyes, but a first order estimate in this case can be made by simply dividing the number of localizations per apical end by 2 as each anti-ALFA nanobody possesses 2 site-specifically bonded Alexa Fluor 647 dyes. This produces the following median numbers of labeled MyoH: 305 for GA-fixed MyoH ALFA, 261 for GA-fixed ALFA-MyoH, 420 for gel-expanded MyoH-ALFA, and 492 for gel-expanded ALFA-MyoH. These estimates are of the same order of magnitude as the MyoH-PAmKate estimate. Thus, it is unlikely that our single-molecule reconstructions are missing a substantial portion of MyoH molecules. The observed distribution of MyoH along the cell axis (Fig. 4E, F) is then likely reflective of the true distribution of the motor as opposed to a labeling artifact, suggesting a true gradient in the number of MyoH molecules distributed along the axis of the conoid with a preference for the conoid tip.

### Gel-Expansion Perturbs MyoH C-Terminal Localization

Comparing the actual sizes of the MyoH structures in GA-fixed and gel-expanded samples requires knowledge of the expansion factor such that reconstructions from different gels can be directly compared. Tubulin imaging in gel-expanded samples provides not only a means of computing an expansion factor but also a common reference for MyoH localizations. Two-color MyoH and tubulin imaging was performed in gel-expanded MyoH-ALFA and ALFA-MyoH samples, with the resulting reconstructions showing improved tubulin labeling (Fig. 5A-D). Apical tubulin localizations increased roughly 3-fold (from 116 to 361, and 86 to 213 for MyoH-ALFA and ALFA-MyoH parasites, respectively) (Fig. S7C, D) and appeared to more closely resemble the expected shape of the conoid. Cross-sections orthogonal to the cell axis showed many more tubulin localizations in close proximity to MyoH localizations (Fig. 5B, D). While the tubulin localization density was increased at all points along the cell axis, the tubulin localization density decreased from the conoid base to the conoid top, which we also observed in GA-fixed samples (Fig. 5E, F).

**Figure 5:**
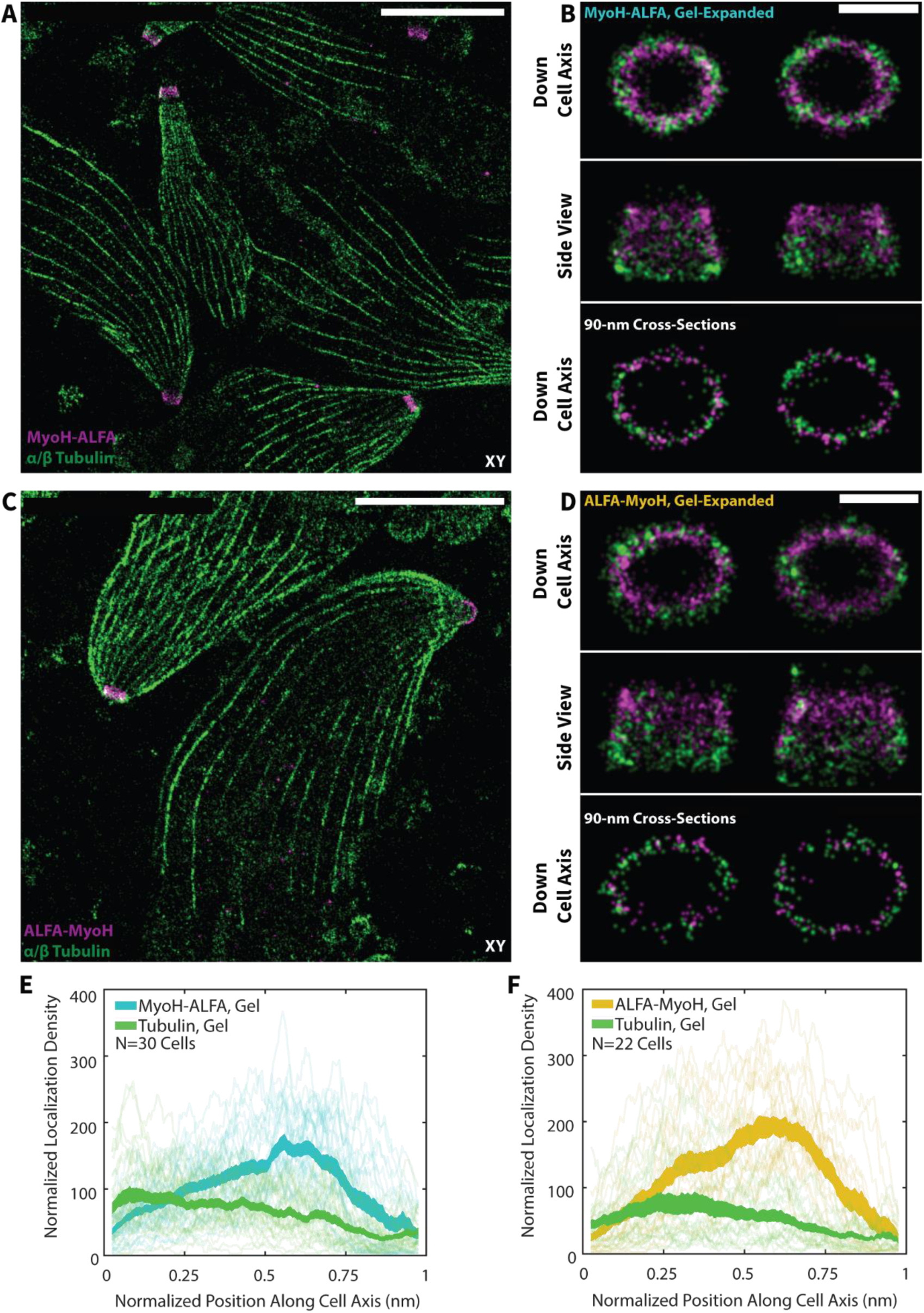
Gel Expansion Improves Conoid Labeling. A) XY projection of 3D localizations from a typical field of view of MyoH-ALFA (magenta) and tubulin (green) labeling in gel-expanded MyoH-ALFA parasites. B) Two aligned, two-color apical reconstructions of gel-expanded MyoH-ALFA (magenta) and tubulin (green) shown in two orientations: down the cell axis (top row) and side view (middle row). Bottom row shows a cross-section 90-nm thick along the cell axis and viewed down the cell axis. C) XY projection of 3D localizations from a typical field of view of ALFA-MyoH (magenta) and tubulin (green) labeling in gel-expanded ALFA-MyoH parasites. D) Two aligned, two-color apical reconstructions of gel-expanded ALFA-MyoH (magenta) and tubulin (green) shown in two orientations: down the cell axis (top row) and orthogonal to the cell axis (middle row). Bottom row shows a cross-section 90-nm thick along the cell axis and viewed down the cell axis. E-F) Normalized localization density vs normalized cell axis for gel-expanded tubulin (green) and MyoH-ALFA (E; N=30 cells) (cyan) or ALFA-MyoH (F; N=22 cells) (yellow). Light lines represent localization density from single cells. Dark lines show the median localization density for all cells with the linewidth representing the standard error of the mean. Scale Bars: (A, C) 5 μm, (B, D) 500 nm.

After correcting for expansion factor using the conoid base as a reference, the radius of N-terminal MyoH localizations from GA-fixed and gel-expanded samples peaked at essentially the same value (medians of 179 nm and 176 nm for GA-fixed and gel-expanded, respectively) (Fig. 6A). Interestingly, the distribution of radii of tubulin and N-terminal MyoH localizations from gel-expanded samples were practically identical (Fig. 6B), differing from the GA-fixed parasites that showed N-terminal MyoH localizations radially further from the cell axis relative to tubulin.

**Figure 6:**
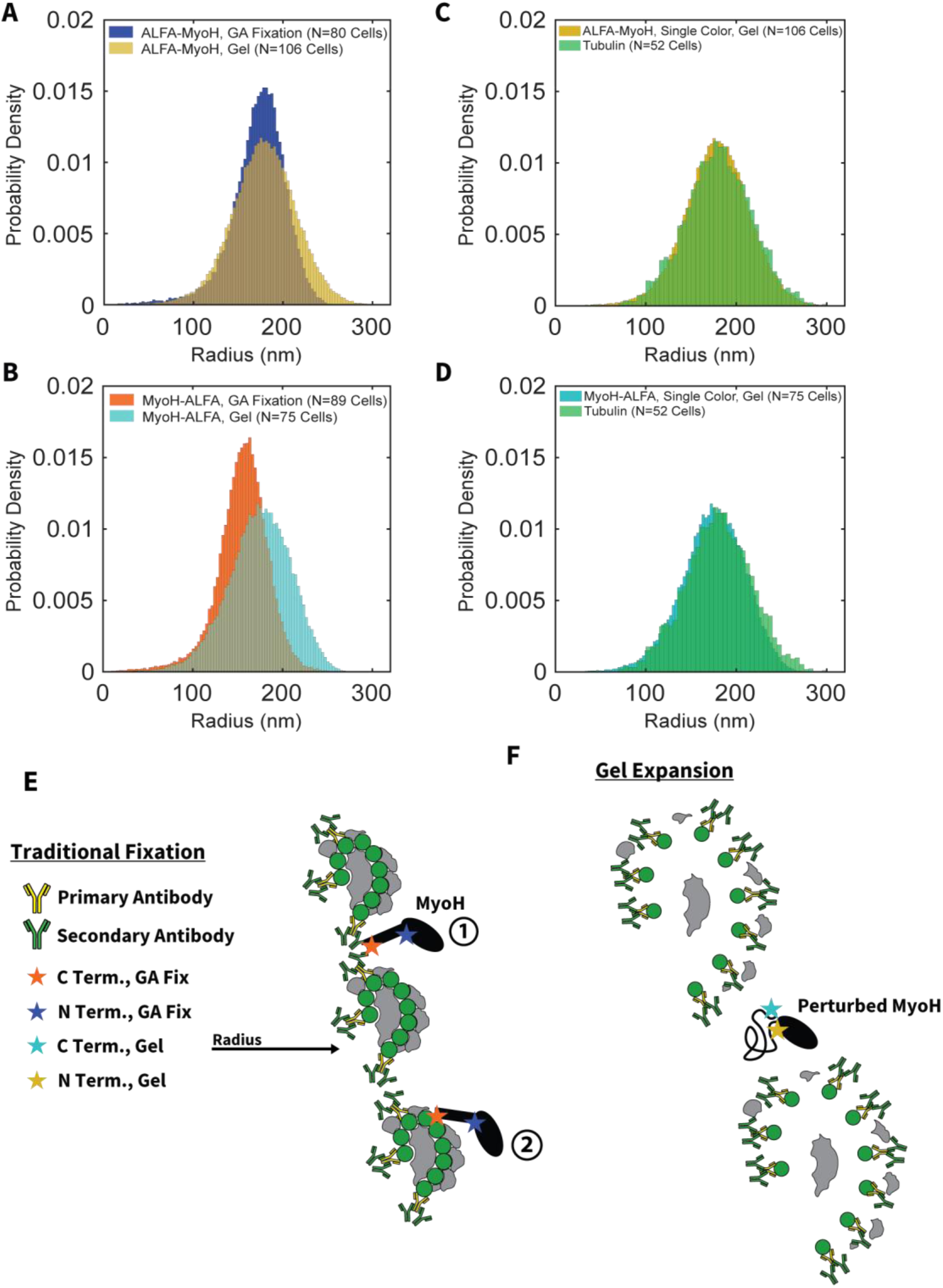
MyoH-ALFA Localizes Differently in Gels Relative to GA Fixed Samples. A) Histogram of radii of ALFA-MyoH localizations from GA fixed samples (N=80 cells) (blue) and gel-expanded samples after correcting for expansion factor (N=106 cells) (yellow). B) Histogram of radii of MyoH-ALFA localizations from GA fixed samples (N=89 cells) (orange) and gel-expanded samples after correcting for expansion factor (N=75 cells) (cyan). C) Histogram of radii of ALFA-MyoH localizations (N=106 cells) (yellow) and tubulin (N=52 cells) (green) from gel-expanded samples after correcting for expansion factor D) Histogram of radii of MyoH-ALFA localizations (N=75 cells) (cyan) and tubulin (N=52 cells) (green) from gel-expanded samples after correcting for expansion factor. E) Models of MyoH organization in GA fixed parasites. Tubulin antibodies preferentially bind the interior face of the conoid fibrils due to greater accessibility. The MyoH C-terminus localizes radially exterior of the tubulin labels. The MyoH N-terminus localizes radially exterior of the C-terminus. Two potential positions of MyoH are consistent with this organization: 1) MyoH inserted between the conoid fibrils and 2) MyoH anchored to the exterior face of the conoid. F) Model of MyoH organization in gel-expanded samples. The MyoH N-terminus localizes similarly in gel-expanded samples as in GA fixed samples. The position of the MyoH C-terminus is perturbed due to the denaturation of the domain during the gel-expansion protocol.

Given that the gel-expanded N-terminal MyoH localizations have similar radii as the GA-fixed localizations, the difference in radii of tubulin localizations seems most likely to result from improved accessibility of the exterior face of the conoid.

In contrast to the N-terminal localizations, C-terminal MyoH localizations in gel-expanded samples had corrected radii peaking at a larger value than in GA-fixed parasites (medians of 176 nm and 157 nm for gel-expanded and GA-fixed, respectively) (Fig. 6C). Additionally, C-terminal MyoH localizations showed a radial distribution nearly identical to that of tubulin in gel-expanded samples (Fig. 6D). Thus, in gel-expanded samples, the MyoH N- and C-termini appear to localize to the same radial position and do not exhibit the radial separation seen in the GA-fixed samples. While one can think of many possible explanations for this result, we believe it is likely caused by perturbation of the MyoH C-terminal localization by gel expansion, as discussed below.

## Discussion

In this study we used 3D single-molecule, super-resolution microscopy to determine the organization of MyoH in the apical complex of *Toxoplasma gondii*. Two separate lines of parasites were created, allowing for labeling of MyoH at either the N- or C-terminus. In both cases, we were able to resolve the ring-shaped profile along the cell axis and the tapered cone profile orthogonal to the cell axis for both conoid-protruded and conoid-retracted parasites. Such reconstructions are expected given previous work establishing the strong association of MyoH with the conoid (25), but our work presents the highest resolution images of MyoH to date in addition to images distinguishing between the N- and C-termini of the protein, i.e., the “head” and “tail” of this novel myosin, respectively. The radial separation observed in GA-fixed parasites thus allowed us to determine that the motor head is on average radially further from the cell axis than the tail. This result represents the first direct measurement of the orientation of MyoH in the apical complex, lending support to previously proposed models of MyoH organization (25).

We wished to add greater context through two-color imaging with tubulin with the goal of localizing MyoH relative to the conoid fibrils. Tubulin labeling of the conoid in GA-fixed parasites was highly heterogenous, failing reconstruct the full conoid and showing heavy bias towards the base of the structure (53). The conoid is comprised of a dense arrangement of proteins (19,20), with several different proteins essentially coating most of the 9 tubulin monomers comprising the protofilaments in the conoid fibrils. Thus, poor conoid labeling is most likely the result of inaccessibility of the conoid tubulin to primary antibodies. Interestingly, the decreasing tubulin localization density along the cell axis suggests that tubulin is more accessible at the conoid base, with the specific proteins surrounding tubulin possibly varying along the cell axis, an idea that has been suggested in the literature (54). Additionally, analysis of the radii of tubulin localizations revealed preferential labeling of the interior face of the conoid fibrils, suggesting that the conoid interior is more accessible to antibodies relative to the exterior. This labeling pattern is consistent with high-resolution models of the conoid fibrils as they show less protein density on the interior face relative to the exterior face (19,20). Thus, tubulin labeling in GA-fixed parasites on its own adds to our understanding of protein organization in the conoid.

Comparing the radial localization of tubulin to MyoH, the 13-nm separation between the C-terminus and conoid tubulin localizations is within the ∼18-nm offset from labeling with primary and secondary antibodies, precluding a precise conclusion as to their relative position but indicating the two are in close physical proximity. The MyoH tail contains 3 ATS1/RCC domains that are suggested to interact with microtubules (55). Additionally, the tail and neck domains of the motor have been shown to be necessary for MyoH localization to the conoid (25), suggesting that the tail and neck of MyoH are anchoring the motor to the conoid. Our results provide key evidence in favor of these hypotheses by showing the nanoscale proximity of the MyoH C-terminus and the conoid tubulin.

Determining an exact position in the conoid architecture for the C-terminus requires knowledge of the arrangement of antibodies labeling the tubulin and the position of the C-terminal amino acid of MyoH in the MyoH structure. Without this information, there are a range of possible MyoH positions that are consistent with our measurements. The C-terminus could be inserted between the conoid fibers as in Model 1 (Fig. 6E). Alternatively, the C-terminus could be anchored more toward the exterior face of the conoid as in Model 2 (Fig. 6E). Homology modeling of the head of MyoH places the N-terminus at the base of the head near the point where the neck begins (56,57). Applying this to our models, the base of the MyoH head then sits either at or beyond the conoid exterior. This is in agreement with previously published models that place the head outside the conoid, allowing the motor to pass the F-actin-glideosome-associated connector (GAC) complex to MyoA in the pellicular space (25,58). In the context of this function, Model 2 would be an advantageous position for the motor to reach the GAC anchored in the parasite membrane and to produce a larger lever arm for force transmission.

We used gel-expansion as a means of increasing epitope accessibility and improving resolution, combining the technique with 3D single-molecule super-resolution microscopy. Gel-expanded reconstructions of both the N- and C-termini of MyoH recovered the same structure as in GA-fixed parasites but with increased localization density across the entire structure. This result highlights the dense environment of the apical complex: the GA-fixed reconstructions were sampled enough to resolve the underlying MyoH structure, yet they were missing several potential localizations. Comparing the number of localizations to estimates of the number of MyoH molecules in the apical end in MyoH-PAmKate, it is likely that the gel-expanded reconstructions are sampling most molecules. If this is indeed the case, the observed apical bias of MyoH along the cell axis is biologically relevant. This could result from the conoid containing an increasing number of binding sites for MyoH from the base to the tip. Alternatively, some aspects of MyoH’s dynamic motor activity could make the protein more likely to unbind closer to the base of the conoid. In either case, this observation highlights the variability in organization of proteins associated with the conoid, which might not be effectively captured by methods that rely on averaging structures along the conoid fibrils.

Gel expansion also improved tubulin labeling, resolving the conoid much more convincingly relative to the GA-fixed samples. In this case, the impact of accessibility is made even clearer. Given our knowledge of the conoid structure and its dimensions, it is immediately obvious that the GA-fixed reconstructions do not recover the expected labeling across the entire conoid, with crowding and local protein density being the likely culprits. However, gel-expanded conoid reconstructions still saw decreasing tubulin labeling density towards the conoid tip, suggesting that our ∼1.7X expansion still left tubulin epitopes inaccessible to primary antibodies. Other approaches such as iterative ultrastructure expansion microscopy (iU-ExM) have recovered relatively complete conoid labeling with expansion factors of 13.9X (59). It is unclear, however, what minimal expansion factor is needed for complete conoid labeling. For our study, the expansion factor was set by the size of the gels in the blinking buffer required for (d)STORM imaging, but the gels prepared can reach an ultimate expansion factor of ∼4 (39). Techniques such as gel re-embedding have been shown to prevent excessive shrinking of polyacrylamide gels, allowing them to maintain an expansion factor of approximately 3 (32,60). 3D (d)STORM of the conoid in re-embedded gels would aid in determining what level of expansion is necessary for complete conoid labeling.

Using tubulin localizations of the conoid base as a means of computing an expansion factor, we rescaled gel-expanded localizations to compare them to GA-fixed localizations. We found that MyoH N-terminal localizations peaked at essentially the same radius with both sample preparations. Additionally, instead of the radial separation between tubulin localizations and N-terminal MyoH localizations observed in GA-fixed samples, gel-expanded samples showed nearly identical distributions of radial coordinates for the N-terminus and tubulin. This result is consistent with labeling of both the interior and exterior face of the conoid, as the additional localizations from the conoid exterior have larger radii and shift the distribution to larger radial values. The C-terminal MyoH localizations had larger radii in gel-expanded samples relative to GA-fixed samples. The distribution of radii was also similar in peak position and shape to that of the N-terminus and tubulin. This result conflicts with the GA-fixed cell model showing the MyoH C-terminus closer to the interior face of the conoid.

This begs the question as to which set of data best represents the ground-truth position of the MyoH C-terminus. If the gel-expanded data are most representative of the ground truth, then both ends of MyoH localize to the same radial coordinate in the apical complex. This would suggest that in the GA-fixed parasites a given MyoH terminus is inaccessible near the other terminus to produce the observed radial separation, requiring a relatively complex model of MyoH organization in the conoid. A simple alternative explanation is that the MyoH C-terminus is altered by gel expansion (Fig. 6F). MyoH itself is a large protein comprised of 1513 amino acids. Given that MyoH has not been resolved by electron microscopy methods that rely on averaging (19,20), it is also likely that the motor has higher mobility relative to the surrounding proteins. The gel expansion protocol involves denaturation at 95 °C in an SDS buffer with high NaCl concentration at pH 9. It is possible that MyoH does not properly refold after this denaturation step, displacing the C-terminus relative to its original position. Additionally, there are on average fewer free amines present in the neck and tail portions of MyoH relative to the head, meaning there are more opportunities for crosslinking into the gel near the N-terminus than the C-terminus, perhaps lending more rigidity to the N-terminus relative to the C-terminus. Thus, while expansion has been shown to have many benefits, especially in the case of the dense protein environment of the *T. gondii* apical complex, we see that gelation and expansion can also introduce perturbations that depend not only on the target protein but also on the specific site at which the protein is labeled.

## Conclusions

In this study we used 3D single-molecule super-resolution microscopy to determine the organization of MyoH in the apical complex of *Toxoplasma gondii*. While the parasite possesses multiple myosin motors, MyoH is unique in that it is necessary for parasite motility, host cell invasion, egress out of host cells, and conoid protrusion. Knowledge of its precise position in the apical complex may aid in further understanding how this motor protein accomplishes these vital roles. While cryo-electron microscopy has provided detailed models of apical complex architecture, MyoH has not been resolved in these views. Our 3D reconstructions in GA-fixed parasites measured the positions of both ends of the motor in the apical complex, resolving a 19-nm separation between the N- and the C-terminus, with the N-terminus radially further away from the cell axis than the C-terminus. This represents the first direct measurement of the orientation of the motor in the apical complex.

Two-color imaging with tubulin in GA-fixed parasites determined that the C-terminus resides close to the interior face of the conoid with the N-terminus located at or beyond the conoid exterior. Gel-expansion improved tubulin labeling and showed N-terminal MyoH localization consistent with GA-fixed data. The MyoH C-terminus, however, localized differently in gels than in GA-fixed samples, showing a radial distribution almost identical to that of the N-terminus. We have attributed perturbation by gel expansion as the most likely cause. These findings have supported aspects of proposed models of MyoH organization, contributed previously unavailable, nanoscale information, and highlighted perturbations to protein localization by gel expansion.

Further work to refine our understanding of MyoH localization in the apical complex could determine the exact sites at which the MyoH tail and neck associate with the conoid fibrils and the precise location of the actin binding pocket of the MyoH head. Dynamic measurements could also build upon the work presented here by investigating the stability of the MyoH association with the conoid and any potential dynamic reorganization of the protein as the conoid protrudes in preparation for host cell invasion.

## Data Availability

Example raw single-molecule blinking data for single-color and two-color datasets for both GA-fixed and gel-expanded samples are available at Ref. (45), in addition to the apical localizations and scripts used for localization and localization processing. Additional data is available upon request.

## Author Contributions

A.B., L.S.-Z., W.E.M, and J.C.B. designed this study. L.S.-Z. performed all cell culture, parasite maintenance, parasite line generation, and fixation. A.B. performed all labeling, gel expansion, imaging, and data analysis. A.E.S.B. contributed scripts for data analysis. A.B. and L.S.-Z. wrote the manuscript with input from all authors.

## Acknowledgements

This work was supported in part by the National Institute of General Medical Sciences Grant No. R35GM118067 (A.B., A.E.S.B, W.E.M.) and the Chan-Zuckerberg Biohub Intercampus Award. L.S.-Z. was supported in part by BARD, the United States–Israel Binational Agricultural Research and Development Fund, Vaadia-BARD Postdoctoral Fellowship Award No. FI-582-2018, the Stanford Maternal and Child Health Research Institute, and Stanford School of Medicine Dean’s Postdoctoral Fellowship. W.E.M. is a Sarafan ChEM-H Institute Fellow. We thank Peter Dahlberg for early work on the MyoH project and for helpful discussions. We thank Ljiljana Milenkovic for guidance and discussions regarding the gel expansion protocol. We thank Michelle Küppers for helpful discussions interpreting data and Melanie Espiritu for help with tissue culture.

