## Supplemental Information for "Organization of Myosin H in the Apical Complex of *Toxoplasma Gondii* Revealed by 3D Single-Molecule Super-Resolution Microscopy"

**Condensed Running Title: Organization of MyoH in *T. gondii***


**Ashwin Balaji,[1, 2] Li-av Segev Zarko,[3, 4, 5] Andrew E.S. Barentine,[2] John C. Boothroyd,[3, 6] W.E. Moerner [1, 2, 6]**

**[1] Biophysics PhD Program, Stanford University, Stanford, CA, 94305, USA**

**[2] Department of Chemistry, Stanford University, Stanford, CA, 94305, USA**

**[3] Department of Microbiology and Immunology, Stanford University, Stanford, CA, 94305, USA**

**[4] Department of Biochemistry and Molecular Biology, University of Texas Medical Branch, Galveston, TX, 77555, USA**

**[5] Sealy Center for Structural Biology & Molecular Biophysics, University of Texas Medical Branch, Galveston, TX, 77555, USA**

**[6] Sarafan ChEM-H, Stanford University, Stanford, CA 94305, USA**

### Supplemental Methods

#### *Western Blot*

Protein samples were prepared from parasite pellets and lysed in SDS sample buffer. Samples were separated by SDS–PAGE and transferred to nitrocellulose membranes. Membranes were blocked in 5% milk in TBST and incubated with a 1:3,500 dilution of mouse monoclonal anti-ALFA antibody (NanoTag, N1582). After washing, membranes were incubated with a 1:3,500 dilution of HRP-conjugated goat anti-mouse IgG1 secondary antibody (Abcam, ab97240). Signal was detected using chemiluminescence.

#### *Blinking Buffer Stock Solutions and Sample Chamber Sealing*

The following stock solutions were used to freshly prepare blinking buffer as needed: 1) 4 kU/ml glucose oxidase (Millipore Sigma, G213), 20 kU/ml catalase (Millipore Sigma, C1345), 25 mM KCl (Fisher Scientific, P217), 4 mM TCEP (Millipore Sigma, 646547), 50% v/v glycerol (Fisher Scientific, BP229) and 22 mM Tris-HCl pH 7.0 (Fisher Scientific, BP1756), stored at −20 °C; 2) 1 M cysteamine-HCl (Millipore Sigma, 30080), stored at −20 °C; 3) 50% w/v glucose (Millipore Sigma, 49139) stored at room temperature; 4) 50 mM NaCl (Fisher Scientific, S271) and 1 M Tris-HCl pH 8.0 (Thermo Fisher Scientific, J22638.AE), stored at room temperature. Samples were then sealed by placing a 22 mm-diameter circular coverslip (Carolina Biological Supply Company, 633035) on top of the chamber and using a Kimwipe to wick away excess moisture at the coverslip edge.

#### *Anti-Tubulin Antibodies Tested on GA Fixed Conoid-Protruded Parasites*

Several sets of primary antibodies were tested for conoid labeling in GA fixed conoid-protruded parasites. These included sheep polyclonal anti-α and anti-β tubulin (Cytoskeleton, Inc., ATN02), mouse monoclonal anti-α tubulin DM1A (Thermo Fisher, 62204) and mouse monoclonal anti-β tubulin 2-28-33 (Millipore Sigma, T5293), mouse monoclonal anti-α tubulin B-5-1-2 (Thermo Fisher, 32-2500) and mouse monoclonal anti-β tubulin 2-28-33 (Millipore Sigma, T5293), mouse monoclonal anti-*T. gondii* α tubulin (Sigma-Aldrich, T5168), and mouse monoclonal anti-α tubulin (ABCD Antibodies, ABCD_AA345) and mouse monoclonal anti-β tubulin (ABCD Antibodies, ABCD_AA344). In all cases, labeling of the conoid tubulin was sparse and heterogenous in GA fixed parasites. We finally selected mouse monoclonal anti-α tubulin B-5-1-2 (Thermo Fisher, 32-2500) and mouse monoclonal anti-β tubulin 2-28-33 (Millipore Sigma, T5293) for fixed cell labeling. The antibodies from ABCD antibodies were selected for gel-expanded samples due to their validated use for tubulin labeling in gel-expanded *T. gondii* (1-3).

***Gel Expansion***

Fixed parasite samples were gel expanded using an adaptation of a previously published protocol (4). Two days before gelation, fresh monomer solution (500 µl 38% Sodium Acrylate (Millipore Sigma, 408220), 250 µl 40% Acrylamide (Bio-Rad, 1610140), 50 µl 2% BIS (Bio-Rad, 1610142), 100 µl 10X PBS) was prepared and stored at -20 °C in 90 µl aliquots.

On the day of gelation, each GA fixed parasite sample was incubated with 1 ml of formaldehyde-acrylamide solution (38 µl 38% Formaldehyde (Millipore Sigma, F8775), 50 µl 40% Acrylamide (Bio-Rad, 1610140), 912 µl PBS) for 5 hours at 37 °C. CellVis chambers were placed in parafilm-sealed humid chambers for incubation to limit evaporation. Humid chambers consisted of a plastic petri dish lined with a damp paper towel with a square of parafilm used to cover the paper towel. When 30 minutes of incubation time remained, 10% TEMED (Bio-Rad, 1610800) in water and 10% ammonium persulfate (Bio-Rad, 1610700) in water were freshly prepared and placed on ice. New humid chambers were then prepared and stored at -20 °C to lower their temperature. 1 aliquot of monomer solution for every 2 gel samples was placed on ice. After the 5-hour incubation, a chilled humid chamber was placed on ice, and two parasite samples were placed in the humid chamber. Formaldehyde/acrylamide solution was removed from the parasite samples, and the parasite samples were very gently blown dry with nitrogen. 5 µl each of 10% TEMED and 10% ammonium persulfate were quickly added to 1 aliquot of monomer solution and then vortexed. 35 µl of this mixture was then placed onto each parasite sample, and a 12-mm diameter circular coverslip (Fisher Scientific, 12-541-001) was then placed on top of each drop of monomer solution. The humid chamber was then closed and allowed to sit on ice for 5 minutes followed by incubation at 37 °C for 1 hour.

The gels were then each covered with 2 ml of denaturation buffer (200 mM SDS (Bio-Rad, 1610418), 200 mM NaCl, 50 mM Tris-Base, pH 9) and placed on an orbital shaker with gentle agitation for 30 minutes. Tweezers were then used to gently remove the top circular coverslip from each gel. Each gel was then gently lifted from the sample chamber with a thin, metal spatula and placed into an Eppendorf tube that was then filled completely with denaturation buffer and incubated in a heat block at 95 °C for 1.5 hours.

Denaturation buffer was then carefully removed from each tube, and each gel was carefully placed into its own beaker filled with 150-200 ml of pure water. After 30 minutes, water in each beaker was replaced with fresh pure water and left to sit covered overnight.

***Gel Imaging Chamber Preparation***

During final antibody incubations, CellVis chambers were first rinsed 3 times with absolute ethanol and then rinsed 3 times with pure water. 0.1 mg/ml poly-L-lysine (Millipore Sigma, P8920) in water was then placed on the coverslip of each chamber and allowed to sit for 10 minutes. Coverslips were then rinsed 3 times with pure water and incubated with a $1:10^6$ dilution of 100 nm 715/755 nm FluoSpheres (Thermo Fisher Scientific, F8799) and 1:12.5 dilution of 100 nm gold nanoparticles (Cytodiagnostics, G-100-20) in PBS for 15 minutes. After rinsing 3 times for 5 minutes with PBS, coverslips were then gently blown dry with nitrogen.

***3D DHPSF Imaging***

3D DHPSF (d)STORM imaging was performed with a previously described, homebuilt, two-color DHSPF microscope (5,6) (Fig. S3) controlled using the PYthon Microscopy Environment (PYME) (7). Briefly, sample illumination was performed with a multimode fiber de-speckler imaged into the sample plane, producing a 35-µm-diameter, circular illumination profile with a flat intensity profile. A quad-pass dichroic mirror separates illumination from fluorescence. A 100X, 1.4NA oil-immersion objective lens collects fluorescence and relays it to a two-channel 4f system. After the first 4f lens, a 660-nm, ultra-flat long-pass dichroic mirror is used to spectrally split the emission. Each resulting arm of the emission pathway contains a double-helix phase mask in the Fourier plane to produce the DHPSF response in each color channel. Finally, a knife-edge prism is placed after each channel's second 4f lens to image each color channel onto opposite quadrants of an EMCCD detector. For all imaging, the focus was set such that fiducials on the coverslip are at the lower edge of the axial range. This focus position is then maintained using a homebuilt focus lock system.

Photoactivation for all imaging was performed with continuous 405 nm illumination. For GA fixed MyoH imaging, photoactivation started on frame 3,000 with 1.2 $W/cm^2$ and increased to the following intensities on the following frames: 2.3 $W/cm^2$ on frame 6,000, 5.8 $W/cm^2$ on frame 10,000, 11.6 $W/cm^2$ on frame 13,000, 23.3 $W/cm^2$ on frame 16,000, 46.6 $W/cm^2$ on frame 19,000, and 64 $W/cm^2$ on frame 21,000. For single-color gel-expanded MyoH samples, photoactivation occurred at the following intensities at the following frames: 0.3 $W/cm^2$ on frame 2,000, 1.2 $W/cm^2$ on frame 4,000, 3.5 $W/cm^2$ on frame 6,000, 7.0 $W/cm^2$ on frame 8,000, 18 $W/cm^2$ on frame 10,000, 35 $W/cm^2$ on frame 12,000, and 64 $W/cm^2$ on frame 14,000. Photoactivation for MyoH in two-color samples began at 0.6 $W/cm^2$ on frame 4000 without any subsequent increase in intensity to limit bleaching of the tubulin label. As the tubulin signal in GA fixed and gel-expanded samples had greater variance relative to the MyoH signal, 405 nm photoactivation was not applied in a uniform manner from field of view to field of view. As localization density decreased, 405 nm intensity gradually increased, starting at 0.6 $W/cm^2$ and peaking at 0.64 $kW/cm^2$.

After cellular imaging, a calibration sample of 100 nm TetraSpeck beads (Invitrogen, T7279) in 1% agarose (5) was used to image the same fiducials in the 3D imaging volume in both color channels simultaneously to generate localizations to compute a registration. Focus was locked to the same z position as used for cellular imaging, and the calibration was laterally scanned to image ~3,000 beads evenly distributed through the imaging volume.

***Localizing and Processing DHPSF Data***

DHPSF data was localized using a Python-based plugin for PYME (8). MyoH images in the 647 nm channel were localized with the following parameters: Detection Threshold: 1.4, Detection Filter Sigma: 5.0. px, ROI Size: 10 px, Background Subtraction: True, Background Range: [-30, 0]. The central slice lobe separation and lobe sigma of the calibration z stack was used for the initial Lobe Separation Guess and Lobe Sigma Guess. Fiducials in 647 nm channel were localized similarly but with a Detection Threshold of 7.0 and without background subtraction. Tubulin images in the 561 nm channel were localized with the following parameters: Detection Threshold: 1.0, Detection Filter Sigma: 5.0. px, ROI Size: 10 px, Background Subtraction: True, Background Range: [-30, 0]. The central slice lobe separation and lobe sigma of the calibration z stack was used for the initial Lobe Separation Guess and Lobe Sigma Guess. Fiducials in the 561 nm channel were localized similarly but with a Detection Threshold of 2.5 and without background subtraction. See Ref. (9) for PYME localization scripts.

Localizations were then assigned z positions using the z stack calibrations, filtered, registered in the case of two-color datasets, drift corrected using smoothed fiducial traces, and merged. Filter parameters for MyoH localizations were as follows: x localization precision < 30 nm, y localization precision < 30 nm, 800 nm < lobe separation < 1600 nm, 140 nm < lobe sigma < 280 nm.  Filter parameters for tubulin localizations were as follows: x localization precision < 50 nm, y localization precision < 50 nm, 700 nm < lobe separation < 1500 nm, 120 nm < lobe sigma < 280 nm. Two-color data was registered using a locally-weighted mean quadratic transformation computed with the TetraSpeck bead localizations from the agarose calibration sample (10). MyoH localizations were merged with a 22-nm radial threshold and a 10-frame temporal window. Tubulin localizations were merged with a 44-nm radial threshold and a 10-frame temporal window. See Ref. (9) for PYME localization processing scripts.

***Estimation of Number of MyoH-PAmKate Molecules per Apical End***

CellVis chambers were first rinsed 3 times with absolute ethanol and then rinsed 3 times with pure water. 0.1 mg/ml poly-L-lysine (Millipore Sigma, P8920) in water was then placed on the coverslip of each chamber and allowed to sit for 10 minutes. Chambers were then rinsed 3X with pure water. Fixed, conoid-protruded MyoH-PAmKate parasite solution was then placed onto the poly-L-lysine-treated coverslips and allowed to sit for 10 minutes before being rinsed 3 times with PBS. Parasites were imaged with the open-aperture psf. 405 nm illumination was first used to photoactivate PAmKate. Low power 561 nm illumination (66 $W/cm^2$) was then used to bring the apical ends into focus and then to measure the total brightness of the apical ends using camera parameters of 13 ms exposures and 284 EM gain. High power 561 nm illumination

(6.6 kW/cm$^2$) was then used to measure the bleaching of the apical ends using camera parameters of 13 ms exposures and 45 EM gain

For each field of view, MyoH-PAmKate data was analyzed first by subtracting dark counts and cropping an 11x11 pixel ROI around each apical center. For each ROI, the background was estimated as the mean of the signal from the edge pixels. After subtracting this background, the apical end signal was estimated as the sum of the central 5 x 5 pixels. The maximum of a 10-frame sliding window mean of the low-power apical signal was used to estimate the total apical brightness. To estimate the single-molecule brightness from the high-power bleaching traces, a change-point algorithm was applied to the apical signal vs time traces from the high-power 561 nm video. Single-molecule brightnesses were pooled together, and the peak of the resulting histogram was used as an estimate of the representative single-molecule brightness. This brightness was then scaled to account for illumination intensity and EM gain differences between the high-power and low-power 561 nm videos. The low-power apical end signal was then divided by the rescaled single-molecule brightness estimate to compute the estimated number of MyoH-PAmKate molecules per apical end.

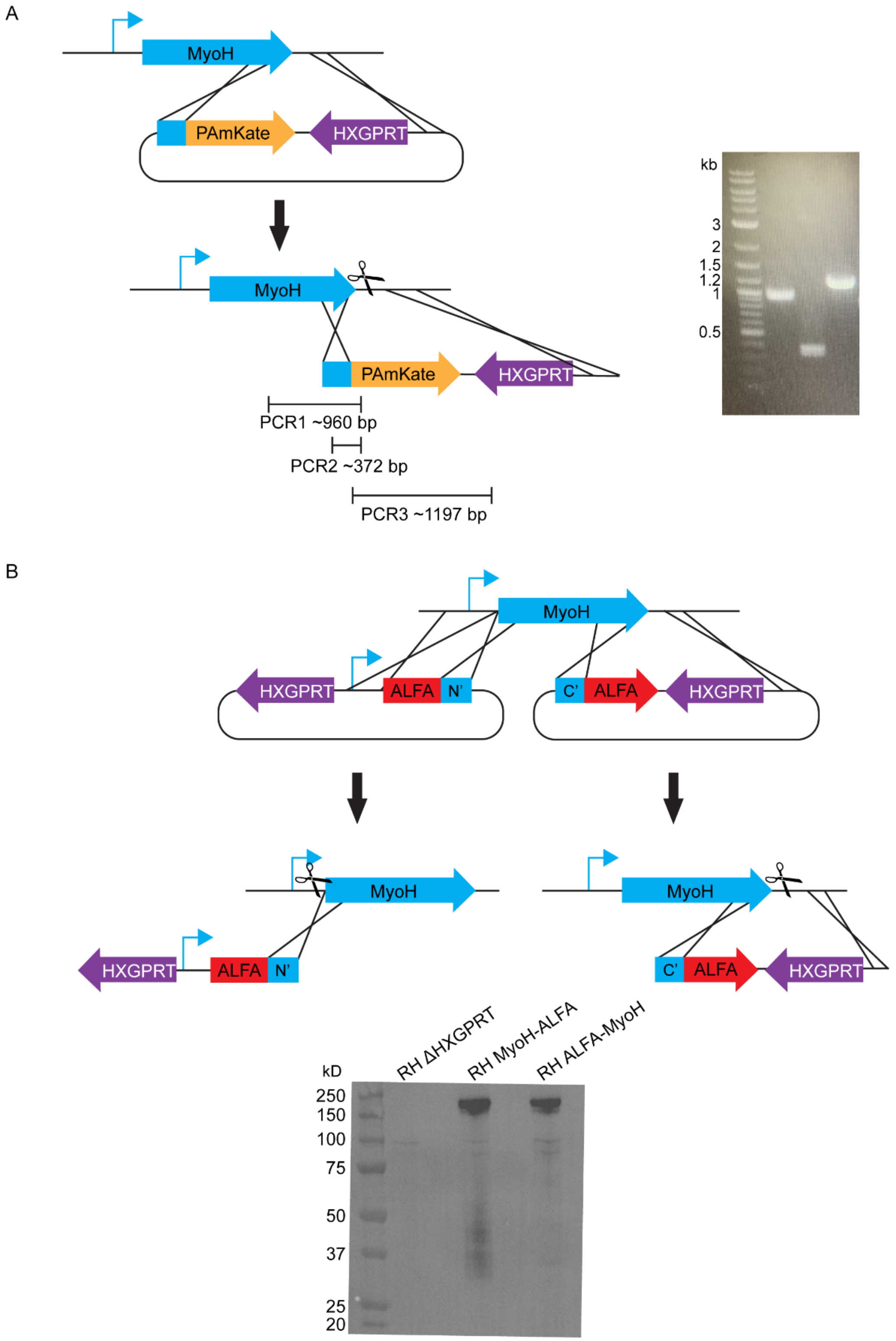


**Figure S1: Endogenous tagging of MyoH and validation of tagged parasite lines** A) Schematic of CRISPR/Cas9-mediated endogenous C-terminal tagging of *MyoH* with PAmKate followed by the HXGPRT selection cassette. The tagging construct, including homology regions, linker, and selection marker, was integrated at the 3’ end of the *MyoH* locus by homologous recombination. Positions of primers used for diagnostic PCR are indicated.

Representative PCR validation of correct integration is shown on the right. B) Strategy for endogenous tagging of *MyoH* at the N-terminus (left) and C-terminus (right) using ALFA tag constructs. For N-terminal tagging, the ALFA tag was inserted upstream of the *MyoH* coding sequence. For C-terminal tagging, the ALFA tag was inserted downstream of the coding sequence. Western blot validation of N- and C-terminally ALFA-tagged MyoH lines is shown on the bottom.

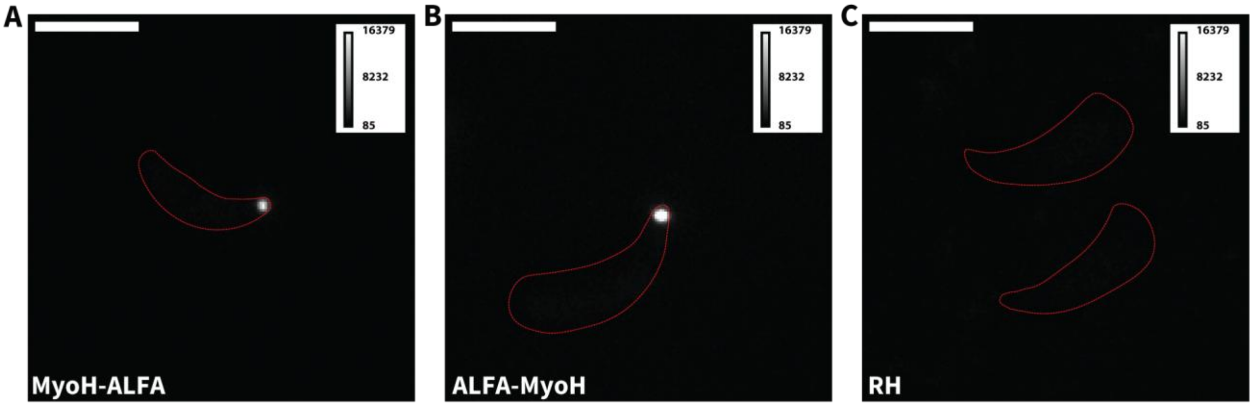


**Figure S2: Specific Labeling of ALFA Tag Lines with anti-ALFA Nanobody** A) Widefield fluorescence image of conoid-protruded MyoH-ALFA parasite labeled with FluoTag X2 anti-ALFA nbAF647. Parasite outlined with dashed red line. Signal can be seen from the apical end of the parasite. B) Widefield fluorescence image of conoid-protruded ALFA-MyoH parasite labeled with FluoTag X2 anti-ALFA nbAF647. Parasite outlined with dashed red line. Signal can be seen from the apical end of the parasite. C) Widefield fluorescence image of conoid-protruded RH parasites labeled with FluoTag X2 anti-ALFA nbAF647. Parasites outlined with dashed red line. No appreciable signal is seen from the parasites.

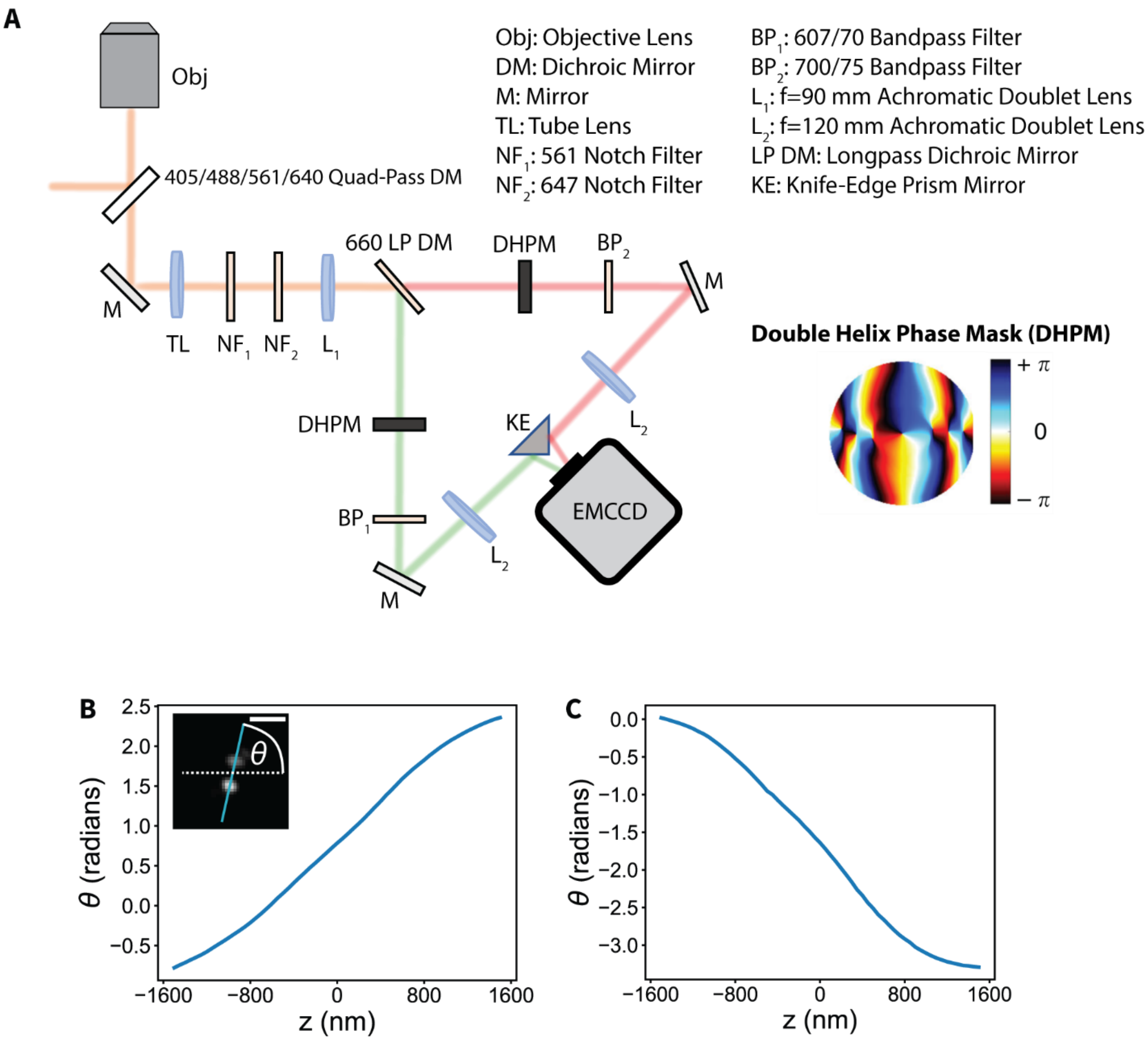


**Figure S3: Homebuilt, Two-Color DHPSF Microscope** A) Schematic of emission pathway of the DHPSF microscope used in this study. Fluorescence from the sample passes through a quad-pass dichroic mirror and is relayed to a two-channel 4f system. A phase mask is placed in the Fourier plane of each arm of the 4f system to produce the DHPSF. B) Lobe angle (θ) vs z calibration curve for the 647 nm channel. Calibration curve produced by scanning a 100-nm diameter Tetraspeck bead through z. Image in plot shows an example frame from the calibration z stack, indicating the horizontal (dashed white line) and the line connecting the DHPSF lobes (cyan). C) Lobe angle vs z calibration curve for the 561 nm channel.

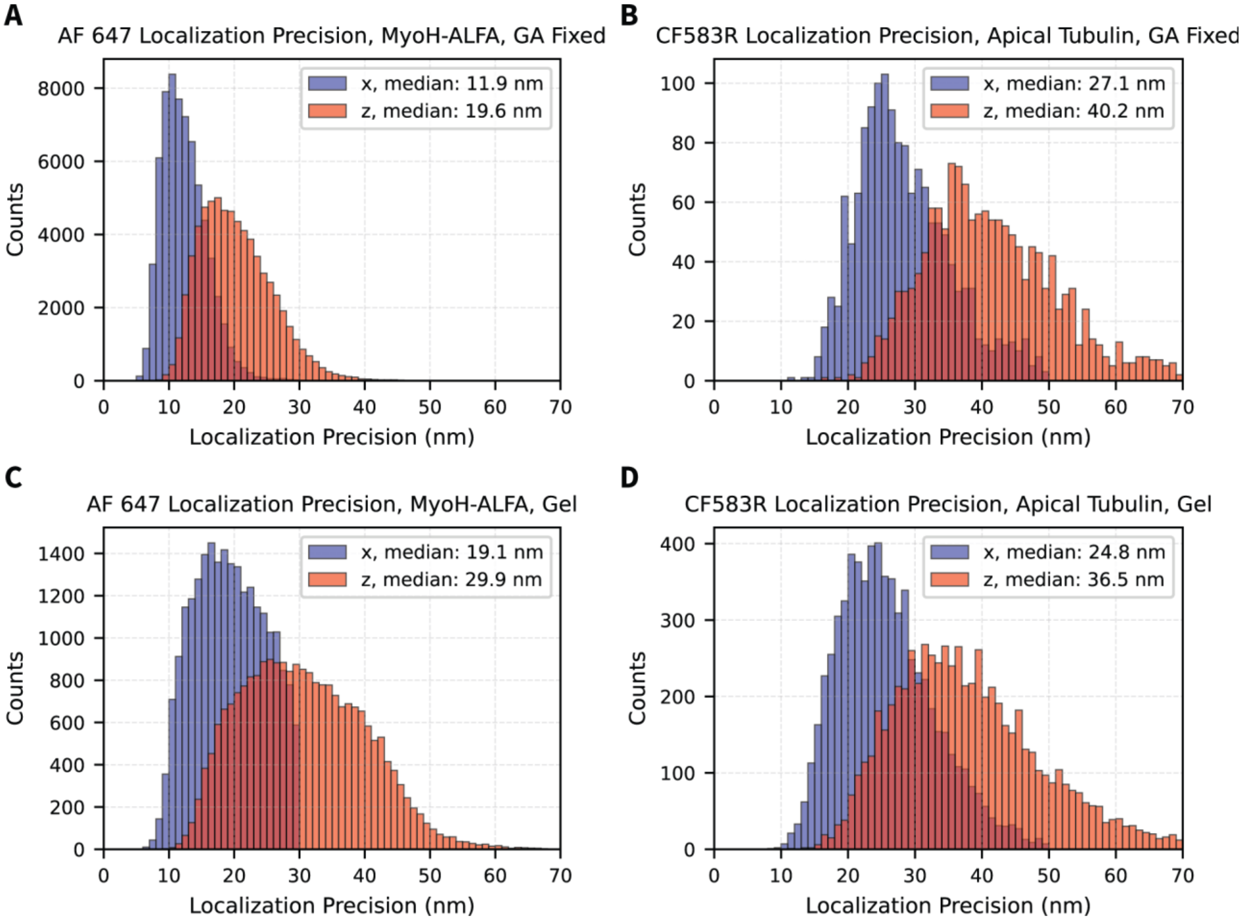


**Figure S4: Localization Precision by Dye and Sample Preparation** A, C) Estimated x (blue) and z (red) localization precision for apical Alexa Fluor 647 localizations (MyoH-ALFA) in GA fixed (A) and gel-expanded (C) conoid-protruded MyoH-ALFA parasites. B, D) Estimated x (blue) and z (red) localization precision for apical CF583R localizations (tubulin) in GA fixed (B) and gel-expanded (D) conoid-protruded MyoH-ALFA parasites.

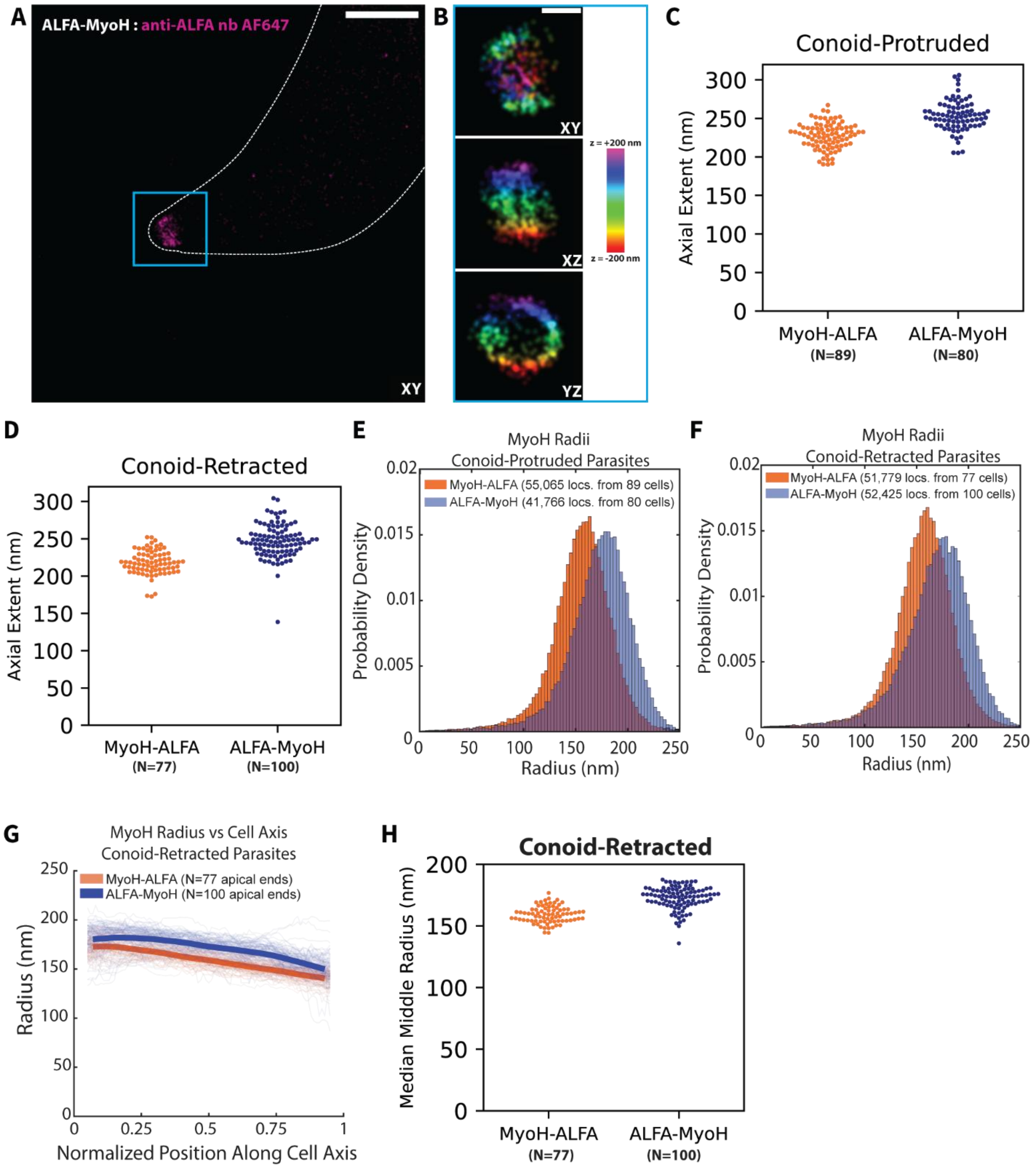


**Figure S5: Single-Color Reconstructions and Radii for GA Fixed, Conoid-Protruded ALFA-MyoH Parasites and for Conoid-Retracted, GA Fixed ALFA-MyoH and MyoH-ALFA Parasites** A) XY projection of localizations from a representative conoid-protruded ALFA-MyoH parasite with the parasite membrane marked with a white, dashed line. A dense set of localizations is seen at the apical tip of the parasite. B) Apical ALFA-MyoH localizations for a representative conoid-protruded parasite viewed in XY, XZ, and YZ projections and colored by z. C) Axial extent of MyoH localizations for each apical end for MyoH-ALFA (orange) (N=89 cells) and ALFA-MyoH (blue) (N=80 cells) from conoid-protruded parasites. D) Axial extent of MyoH localizations for each apical end for MyoH-ALFA (orange) (N=77 cells) and ALFA-MyoH (blue) (N=100 cells) from conoid-retracted parasites. E) Histogram of radii of MyoH-ALFA localizations (N=55,065 localizations from 89 cells) (orange) and ALFA-MyoH localizations (41,766 localizations from 80 cells) (blue) both for conoid-protruded parasites. F) Histogram of radii of MyoH-ALFA localizations (N=51,779 localizations from 77 cells) (orange) and ALFA-MyoH localizations (52,425 localizations from 100 cells) (blue) both for conoid-retracted parasites. G) Radius vs normalized position along cell axis for MyoH-ALFA (orange) (N=77 cells) and ALFA-MyoH (blue) (N=100 cells) localizations from conoid-retracted parasites. Light lines represent binned radial values for single cells. Dark lines represent the median radius for all cells with the linewidth representing the standard error of the mean. H) Median middle radius of each apical end for MyoH-ALFA (orange) (N=77 cells) and ALFA-MyoH (blue) (N=100 cells) from conoid-retracted parasites. Each dot shows the median radius of localizations ranging from 0.45 to 0.55 along the normalized cell axis for a single cell. Scale Bars: (A) 1 µm, (B) 200 nm.

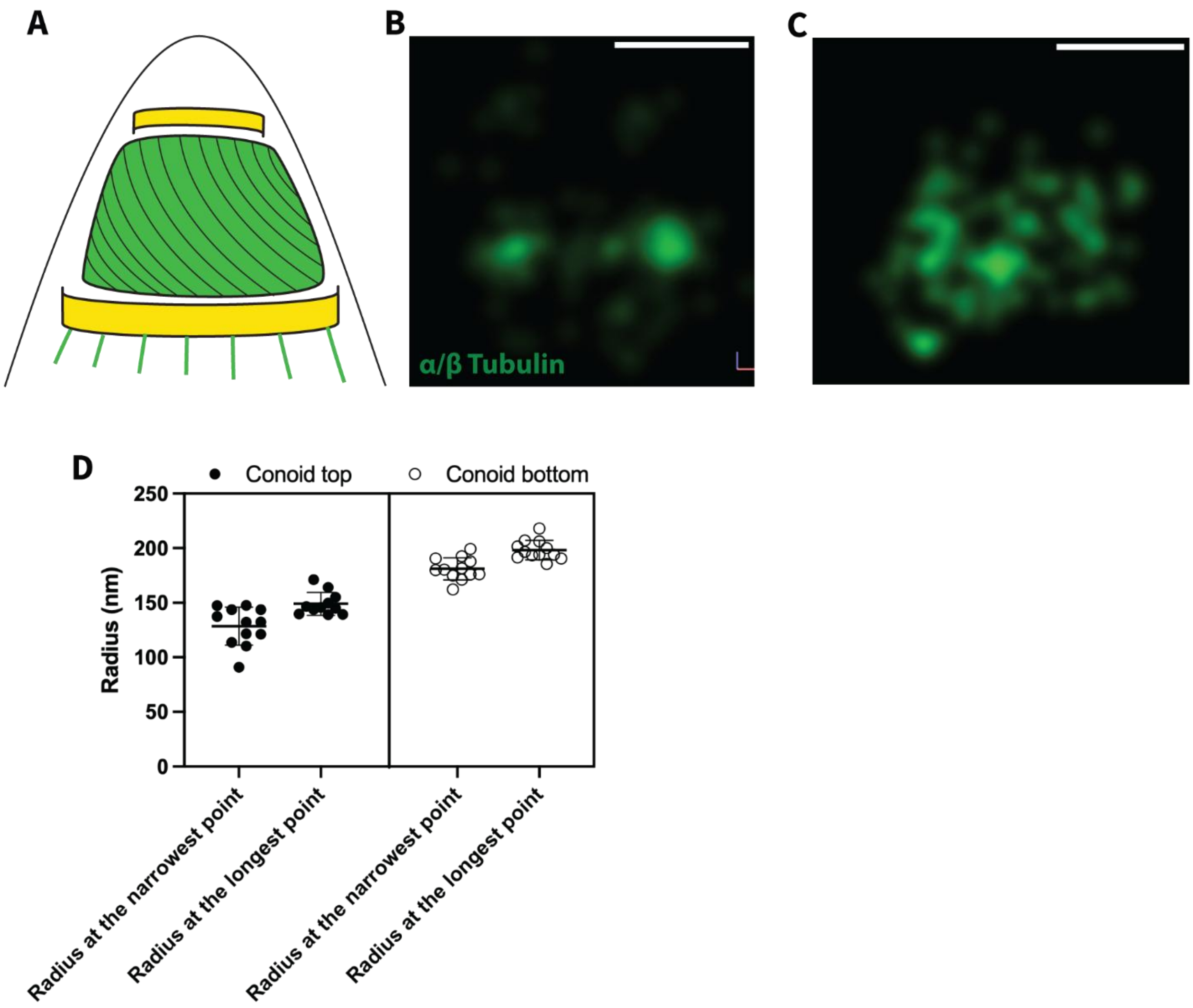


**Figure S6: Conoid Tubulin Labeling is Highly Heterogeneous in GA Fixed Parasites** A) Cartoon of the T. gondii apical complex showing the side profile of the conoid in green. B) Side profile of tubulin localizations from the apical end of a GA fixed, conoid-protruded MyoH-ALFA parasite. Labeling is sparse in this case with heavy bias for the base of the conoid. C) Side profile of tubulin localizations from the apical end of a GA fixed, conoid-protruded MyoH-ALFA parasite. In this case the rough shape of the conoid can be seen. D) Quantification of conoid radius measured from cryo-ET data (11) of conoid-protruded parasites at the conoid top and bottom. Because the conoid in the tomograms does not exhibit circular symmetry, radii were measured along the short and long axes. Each point represents an individual conoid. Scale Bars: (B, C) 200 nm.

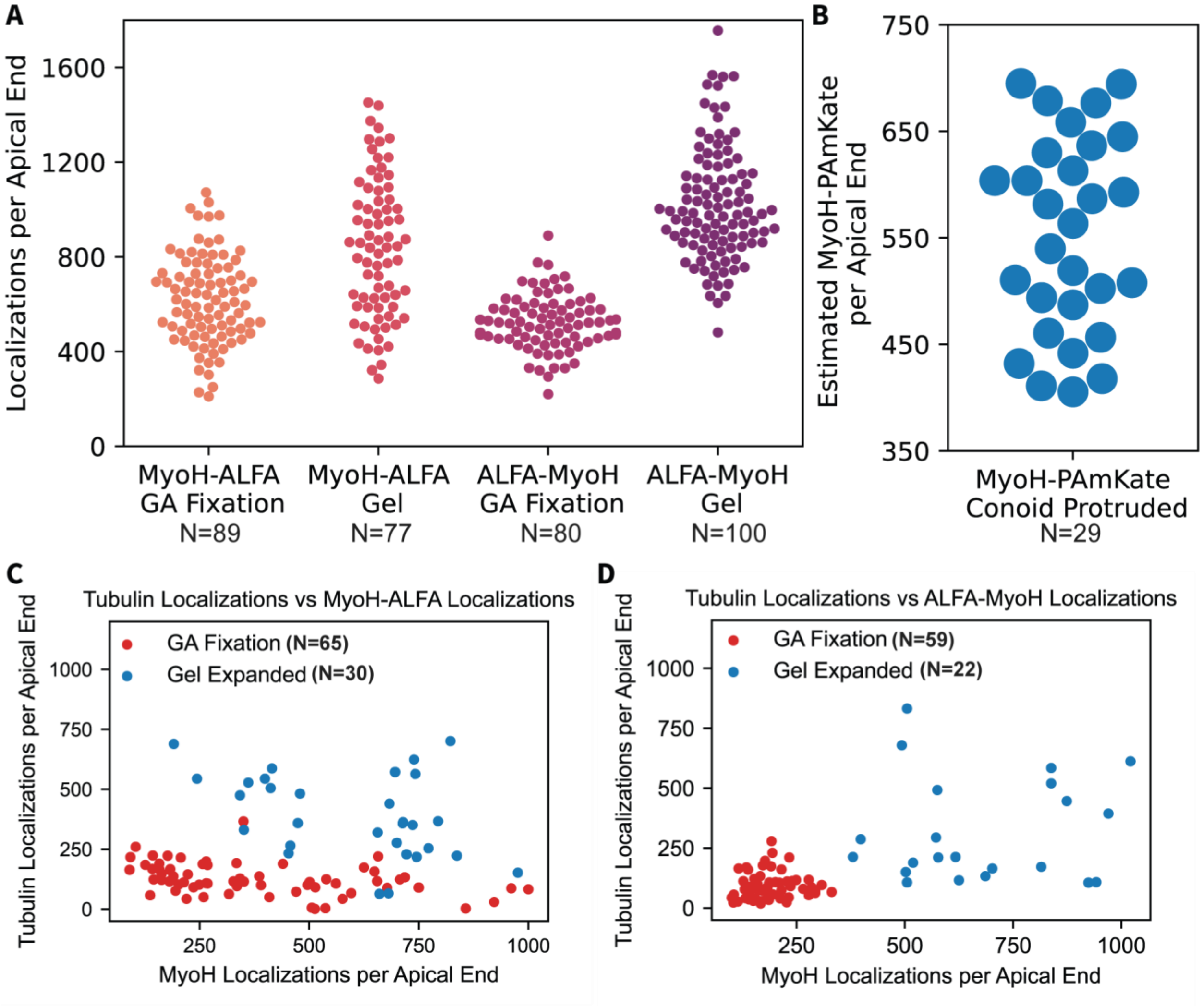


**Figure S7: Number of MyoH Localizations Per Apical End** A) Number of MyoH localizations per apical end in GA fixed and gel-expanded conoid-protruded parasites. B) Estimated number of MyoH-PAmKate molecules per apical end from bleaching experiments in conoid-protruded MyoH-PAmKate parasites. C) Number of tubulin and MyoH-ALFA localizations in GA fixed and gel-expanded conoid-protruded parasites. D) Number of tubulin and ALFA-MyoH localizations in GA fixed and gel-expanded conoid-protruded parasites.

### Table S1: Primers and Oligonucleotides

| Number | Sequence (5'→3') | Description |
|---|---|---|
| 20 | AGCGGAAAGTGGACGGCATTGGTAGTGGGAGCAACGGCAGCAGCGGATCCatgagctcgagatctatgagc | PAmKate amplification and cloning into pSAG1:U6-Cas9:sgUPRT |
| 21 | GTGTGTTTCCTTTGTCGATTTGAGAAGTGAGCACAACGGTGATTAATTAAattaagcttgtgccccagttt | PAmKate amplification and cloning into pSAG1:U6-Cas9:sgUPRT |
| 24 | TCTAGAACTAGTGGATCCCCCtccaccgcgatgagctcgagatctatgag | PAmKate and SAG1 3' UTR amplification and cloning into pGRA1-HXGPRT-3xHA |
| 25 | gacaagtgttctggcaggctacagtgacacCGCTCTAGAACTAGTGGATC | PAmKate and SAG1 3' UTR amplification and cloning into pGRA1-HXGPRT-3xHA |
| 42 | ctccctagcaaactggggcacaagcttaattaaTTAATTAATCACCGTTGTGCTCACTTCTCA | Adding a stop codon downstream of PAmKate |
| 43 | TGAGAAGTGAGCACAACGGTGATTAATTAAttaattaagcttgtgccccagtttgctagggag | Adding a stop codon downstream of PAmKate |
| 64 | ggaggaggaagtggaggaggaagtatgagctcgagatctatgagcgagct | Adding a linker sequence (ggaggaggaagt)x2 upstream of PAmKate |
| 65 | GAGGAATCGAGTGGATGCGCCCTCCGTGGT | Adding a linker sequence (ggaggaggaagt)x2 upstream of PAmKate |
| 66 | TCTAGAACTAGTGGATCCCCCtccaccgcgTTCTTTGGGCGTGGAATTTCTT | MyoH C-terminus amplification and cloning into pGRA1-PAmKate-HXGPRT |
| 67 | gctcatacttcctcctccacttcctcctccGTTGTAGGCCATGGGATCCCAGTTCGGATTCACCGACGGACG | MyoH C-terminus amplification and cloning into pGRA1-PAmKate-HXGPRT |
| 68 | Cgggcgggtttgaatgcaaggtttcgtgctggatctttttggcggctgcc | MyoH 3′ UTR amplification and cloning into pGRA1-PAmKate-HXGPRT |
| 69 | AGCACCAGTACTACAGCCTTCGAAGCTTGATCGCTAGGGAGCAGAGAGAG | MyoH 3′ UTR amplification and cloning into pGRA1-PAmKate-HXGPRT |
| 169 | CCGAGCAGGCTGGAGGAGGAGCTGAGGAGGAGGCTGACCGAGtaaTTAATTAATCACCGTTGTGCTCACTTC | Replacing PAmKate with ALFA tag |
| 170 | CTCGGTCAGCCTCCTCCTCAGCTCCTCCTCCAGCCTGCTCGGGTTGTAGGCCATGGGATCCCAGTTCGGATT | Replacing PAmKate with ALFA tag |
| 127 | cgctagggagcagagagagtc | Amplifying the PAmKate or ALFA tag cassette |
| 234 | TTCTTTGGGCGTGGAATTTCTTCG | Amplifying the PAmKate or ALFA tag cassette |
| 217 | atctttttggGTTTTAGAGCTAGAAATAGC | Cloning sgMyoH_3′UTR into pSAG1:U6-Cas9:sgUPRT |
| 218 | cttagttgtaAACTTGACATCCCCATTTAC | Cloning sgMyoH_3′UTR into pSAG1:U6-Cas9:sgUPRT |
| 121 | GCCAGTCGATTTGAGACGCG | MyoH-PAmKate PCR verification 1 |
| 125 | ccgtacatgaagctggtagcca | MyoH-PAmKate PCR verification 1 and 2 |
| 124 | GTCTCCACTCTTCTCAGCTGTGTT | MyoH-PAmKate PCR verification 2 |
| 14 | atgagctcgagatctatgag | MyoH-PAmKate PCR verification 3 |
| 29 | atcgagccttgagcgtttct | MyoH-PAmKate PCR verification 1 |

| Number | Sequence (5'→3') | Description |
|---|---|---|
| 130 | TAGTTCCAGTCGACGGATCCAC | Generate reduced version of pGRA1-HXGPRT-3xHA (pGRA1-HXGPRT-R) |
| 131 | AAGCTTGATcagcacgaaaccttg | Generate reduced version of pGRA1-HXGPRT-3xHA (pGRA1-HXGPRT-R) |
| 261 | TGAATGCAAGGTTTCGTGCTGATCAAGCTTGTCACCCAGCATGGACTGTG | MyoH promoter and N-terminus amplification and cloning into pGRA1-HXGPRT-R |
| 262 | CTCTAGAACTAGTGGATCCGTCGACTGGAACTATCGTTAGCGTGAGGGAGTTTGG | MyoH promoter and N-terminus amplification and cloning into pGRA1-HXGPRT-R |
| 173 | CTGAGGAGGAGGCTGACCGAGCCGCCAAAGAAGGC | Adding ALFA tag to MyoH N-terminus |
| 174 | CTCCTCCTCCAGCCTGCTCATTTTGAGTCGAAAAAGGAATC | Adding ALFA tag to MyoH N-terminus |
| 271 | ACCGCGGTGTCACTG | Amplifying the ALFA tag cassette |
| 274 | TCGTTAGCGTGAGGGAGTTTG | Amplifying the ALFA tag cassette |
| 228 | CCGCCAAAGAGTTTTAGAGCTAGAAATAGC | Cloning sgMyoH_5′UTR into pSAG1:U6-Cas9:sgUPRT |
| 229 | CATTTTGAGTCAACTTGACATCCCCATTTAC | Cloning sgMyoH_5′UTR into pSAG1:U6-Cas9:sgUPRT |

### Table S2: Plasmids

| Plasmid Name | Source |
|---|---|
| pGRA1-HPT-3xHA | Ref. (12) |
| pSAG1:U6-Cas9:sgUPRT | Ref. (13) |
| pXyl_PAmKate_PopZ | Ref. (14) |
| pGRA1-PAmKate-HPT | This study |
| pGRA1-MyoH-PAmKate-HPT | This study |
| pGRA1-MyoH-ALFA-HPT | This study |
| pGRA1-HPT-R | This study |
| pGRA1-HPT-R-ALFA-MyoH | This study |
| pSAG1:U6-Cas9:sgMyoH_3’UTR | This study |
| pSAG1:U6-Cas9:sgMyoH_5’UTR | This study |
